# Genetic context controls early microglia-synaptic interactions in mouse models of Alzheimer’s disease

**DOI:** 10.1101/2023.04.28.538728

**Authors:** Sarah E. Heuer, Kelly J. Keezer, Amanda A. Hewes, Kristen D. Onos, Kourtney C. Graham, Gareth R. Howell, Erik B. Bloss

**Affiliations:** The Jackson Laboratory, Bar Harbor, ME 04609, USA; Tufts University Graduate School of Biomedical Sciences, Boston, MA 02111, USA; Graduate School of Biomedical Sciences and Engineering, University of Maine, Orono, Maine 04469, USA

## Abstract

Common features of Alzheimer’s disease (AD) include amyloid pathology, microglia activation and synaptic dysfunction, however, the causal relationships amongst them remains unclear. Further, human data suggest susceptibility and resilience to AD neuropathology is controlled by genetic context, a factor underexplored in mouse models. To this end, we leveraged viral strategies to label an AD-vulnerable neuronal circuit in CA1 dendrites projecting to the frontal cortex in genetically diverse C57BL/6J (B6) and PWK/PhJ (PWK) *APP/PS1* mouse strains and used PLX5622 to non-invasively deplete brain microglia. Reconstructions of labeled neurons revealed microglia-dependent changes in dendritic spine density and morphology in B6 wild-type (WT) and *APP/PS1* yet a marked stability of spines across PWK mice. We further showed that synaptic changes depend on direct microglia-dendrite interactions in B6.*APP/PS1* but not PWK.*APP/PS1* mice. Collectively, these results demonstrate that microglia-dependent synaptic alterations in a specific AD-vulnerable projection pathway are differentially controlled by genetic context.

## INTRODUCTION

Alzheimer’s disease (AD) is a neurodegenerative disorder in which individuals exhibit decline across multiple cognitive domains including learning and memory, and executive functioning^1^. AD is defined by the accumulation of amyloid beta (Aβ) plaques and neurofibrillary tangles in the brain^2^. These are associated with progressive yet selective patterns of synaptic disruption that emerge in entorhinal and hippocampal neural circuits before becoming widespread across frontal and parietal association areas^3, 4^. Selective vulnerability of these circuits occurs alongside robust neuroinflammation^5^. A plausible hypothesis is that activation of microglia in response to Aβ plaque deposition causes neural circuit disruption which is supported by the demonstration that microglia regulate circuit connectivity during development and in the healthy, adult brain^6, 7^. Further, microglia depletion in AD mouse models has been shown to reduce late-stage synapse loss on hippocampal neurons^8, 9^. However, these experiments are confounded by the observation that microglia depletion caused spatial redistribution of parenchymal Aβ plaques to the cerebrovasculature^10, 11^. Thus, despite much research, the precise relationship between Aβ deposition, microglia activation, and synaptic connectivity among neurons remains unclear.

Age at symptom onset is variable among AD patients carrying similar rare high-risk mutations in *APP* and *PSEN1* and bearing similar plaque loads, implicating genetic context as an important factor controlling AD progression^12^. Cognitive resilience has also been observed in subsets of late onset AD (LOAD) patients, suggesting additional genetic components shape the cellular events mediating cognitive decline^13, 14^. However, previous studies of synaptic dysfunction using AD mouse models have been performed almost exclusively on the inbred C57BL/6J (B6) genetic background. Incorporation of genetic diversity into AD mouse models has become better appreciated through studies of traditional transgenic AD mouse models on genetically diverse mouse strains^15–19^. We have shown that despite identical patterns of Aβ plaque deposition, wild-derived PWK/PhJ (PWK) mice carrying the *APP/PS1* transgene (PWK.*APP/PS1*) exhibit cognitive resilience compared to traditionally-studied B6.*APP/PS1* inbred mice^18^. Intriguingly, PWK and PWK.*APP/PS1* mice also contain different proportions of transcriptionally-defined microglia states compared to B6, suggesting microglia may be a potential factor mediating resilience to brain Aβ deposition^19^.

Here, we examined how genetic context influences the role of microglia on synaptic changes during early stages of Aβ plaque deposition. We used a viral approach^20^ to gain genetic access to an AD-vulnerable neuronal circuit^21^ that connects hippocampal area CA1 to the prefrontal cortex (PFC) in B6 and PWK wild-type (WT) and *APP/PS1* transgenic (TG) mice, permitting rigorous comparisons across equivalent neuronal populations. The CSF1R inhibitor PLX5622 was used to deplete microglia while keeping Aβ plaque deposition unaltered. Dendritic spine density (a proxy for synaptic number) and spine morphology (a proxy for synaptic stability, plasticity and strength^22–24)^ was quantified across the dendritic compartments of CA1 pyramidal cells. We found a microglia-dependent increase in spine density on proximal oblique dendrites from B6.*APP/PS1* mice that was accompanied by a shift towards smaller spines; both effects were completely absent in PWK.*APP/PS1* mice. Further supporting a context-dependent role for microglia in synapse plasticity during AD, B6.*APP/PS1* but not PWK.*APP/PS1* mice showed differential remodeling of synapses on branches directly contacted by microglia processes. Collectively, these results provide strong evidence that the mechanisms driving synaptic responses to amyloid depend on genetic context.

## RESULTS

### CSF1R inhibition depletes microglia without altering plaque pathology in female *APP/PS1* mice

To determine how microglia influence CA1 neurons we formulated the CSF1R inhibitor PLX5622 in mouse diet as described previously^10^. A 3-week pilot study was performed to determine the safety and efficacy of PLX5622 diet in adult (2.5m) female PWK compared to B6 mice. Flow cytometric analysis of isolated myeloid cells from brain hemispheres and immunofluorescence analysis of dorsal CA1 found that PLX5622 diet significantly depleted microglia by 50% in B6, and 80% in PWK (**Figure S1A-C**). No within-strain differences in body weights or quantity of diet consumption were observed between PLX5622 and control diet groups (**Table S1**). Flow cytometric analysis of peripheral blood revealed no effects on composition of major immune cell populations (**Figure S1D-M**).

To test how genetic context controls amyloid and microglia to CA1 circuit vulnerability, we generated cohorts of B6.*APP/PS1* and PWK.*APP/PS1* TG female mice and WT littermate counterparts (n=12 per strain/genotype group). At 3 months of age, we performed dual intracranial injections of recombinant adenoassociated virus (AAVretro-Cre in prefrontal cortex (PFC) and FLEX-*rev*-EGFP in CA1, see **Methods**) to drive EGFP expression in CA1 neurons that project to the PFC. At 4 months of age (when *APP/PS1* plaque deposition is first observable^25^), we placed 6 mice per strain/genotype group on PLX5622 diet and left 6 mice/group on Purina 5K52 control diet for 4 months until all mice reached 8 months of age (**Figure 1A**). Mice were perfused with fixative, coronally sectioned and immunolabeled for markers of amyloid pathology (X34) and microglia (IBA1). The laminar structure in CA1 reflects distinct afferent pathways, so data were analyzed across subregions: stratum lacunosum moleculare (SLM), stratum radiatum (SR), and stratum oriens (SO) (statistics for each region reported in **Table S2)**.

**Figure 1:**
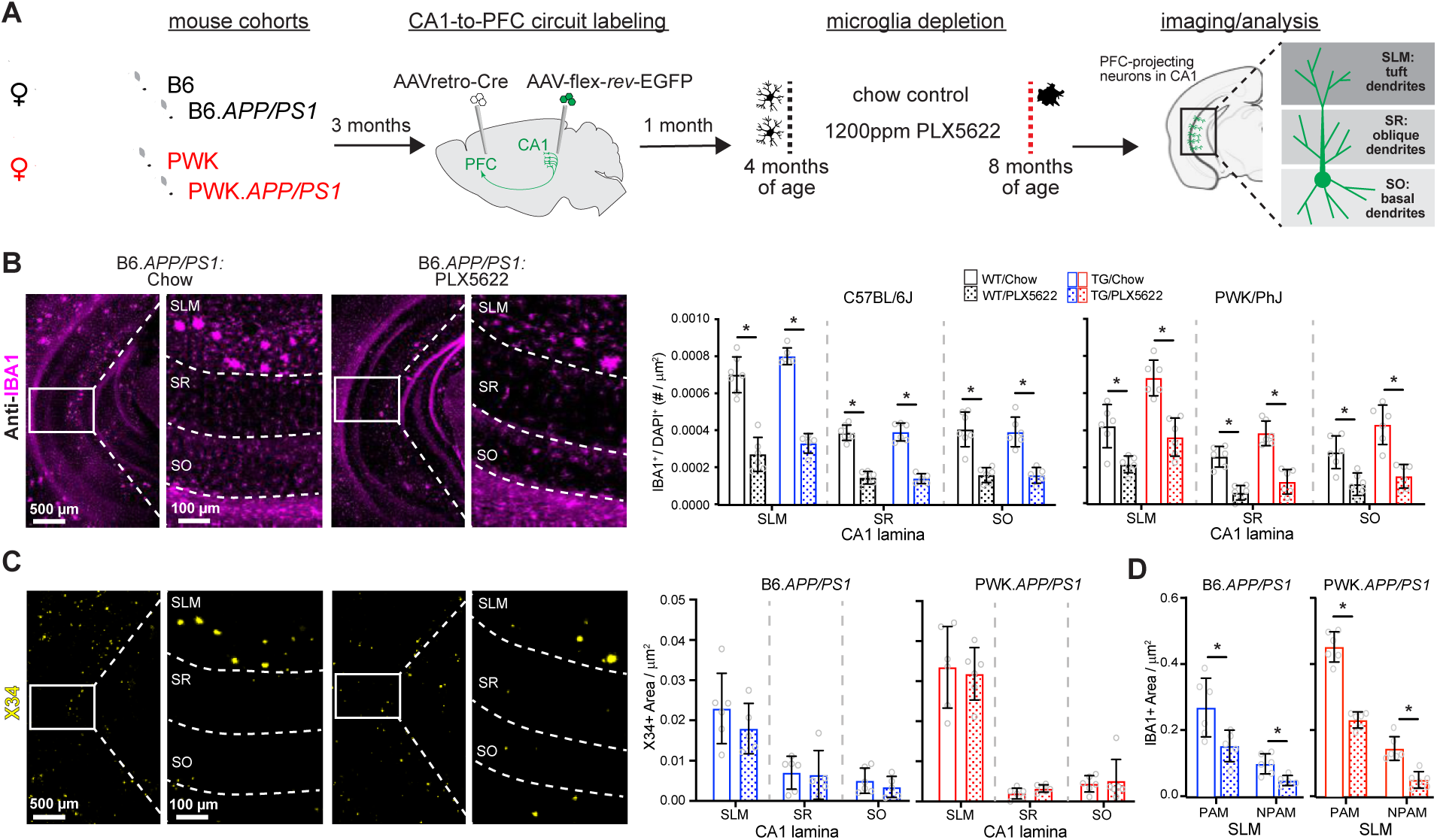
Evaluation of microglia composition and amyloid pathology across CA1 in WT and *APP/PS1* mice. **(A)** Experimental outline (see **Methods** for additional details). **(B)** IBA1+ microglia from B6.*APP/PS1* TG control and PLX5622 mice (left). Quantification of IBA1+/DAPI+ microglia across CA1 lamina (right). Datapoints represent individual mice; error bars are ± SD; asterisks denote comparisons (p<0.05) identified between control and PLX5622 groups (right) after corrections for multiple comparisons. SLM, stratum lacunosum moleculare; SR, stratum radiatum; SO, stratum oriens. **(C)** X34+ Aβ plaques in B6.*APP/PS1* TG control and PLX5622 mice (left). Quantification of X34+ Aβ plaque area across CA1 lamina (right), plotted as described above. **(D)** Quantification of IBA1+ area from SLM defined as plaque-associated (PAM) or non-plaque associated microglia (NPAM). Points represent mean values calculated for individual mice and analyzed with two-tailed nonparametric t-tests. Statistical analyses performed on B6 and PWK separately. For (B)-(C) *adjusted p<0.05 Bonferroni post-hoc tests. For (D) *p<0.05 nonparametric two-tailed t-test (**Table S2**).

With control diet, the density of microglia was several-fold higher in SLM compared to SR or SO. Both strains, regardless of *APP/PS1* genotype, showed significant PLX5622-mediated reductions of microglia (**Figure 1B, Figure S1N**). Depletion efficiencies were dependent on laminae, with greater microglia depletion in SO or SR compared to SLM. We examined X34+ Aβ plaque pathology and found that plaque density varied across CA1 laminae but did not differ between PLX5622 and control diet animals of each strain (**Figure 1C, Figure S1N**). Unlike previous reports^10, 11^, PLX5622 treatment did not result in increased cerebral amyloid angiopathy (CAA)^26^ (**Figure S1O**). We also compared plaque-associated microglia (PAM, defined as IBA1+ microglia localized within 100μm diameter circle from plaque center) to non-plaque associated microglia (NPAM) in SLM and found that both PAM and NPAM were significantly depleted (**Figure 1D, Figure S1P**). This approach allowed for specific evaluation of microglia-neuron interactions without confounding changes to Aβ plaque pathology.

### Strain-specific effects of microglia depletion on proximal dendritic synapses of CA1-to-PFC projection neurons

The vast majority of excitatory synaptic inputs to CA1 pyramidal cells are made onto proximal oblique or basal dendrites^27, 28^. Since these dendritic compartments are close to the site of action potential generation, synapses formed onto these branches strongly affect the output patterns of the cell. Previous work has highlighted structural remodeling of proximal dendrites in response to Aβ pathology, an effect that changes the integrative properties of the dendrites^29^. We first examined dendritic structure in SR via three-dimensional reconstructions of CA1-to-PFC projection neurons and found no differences among dendritic lengths or branch points across B6 mice regardless of genotype or treatment, but a significant increase in dendrite length between PWK WT and PWK TG PLX5622 mice (**Figure 2A**). While Sholl analyses revealed no significant change in dendritic length at specific distances from the soma in B6 mice, both treatment and genotype effects were present in PWK (**Table S3**) such that lengths were increased in PWK TG PLX5622 compared to PWK WT control mice at several distances from the soma (**Figure S2A**). Virtually all excitatory synapses are formed at spines on CA1 pyramidal cells, and approximately all spines contain a single excitatory synapse^30, 31^. Previous studies across AD mouse models have found varying degrees of non-specific synaptic loss in the hippocampus^9, 32, 33^. To elucidate the circuit specificity of Aβ- and microglia-dependent synaptic changes, we imaged SR oblique dendrites from EGFP-labeled CA1-to-PFC projection neurons and reconstructed dendritic spines, calculating spine densities (spines/μm) for each segment (**Figure 2B**). Our measured densities were comparable to those found with array tomography (AT) and by serial section electron microscopy (ssEM) on the same branch types from mouse CA1^27, 34^ (**Figure S2B**). Our analysis revealed significant main and interactive effects from dendrites of B6 mice with B6 TG control branches having significantly higher spine densities than branches from B6 WT control and WT PLX5622. This effect was absent in B6 TG PLX5622-treated mice (**Figure 2C**), suggesting Aβ-dependent increases in spine densities required microglia. None of these effects were evident on oblique branches from PWK mice regardless of genotype and treatment.

**Figure 2:**
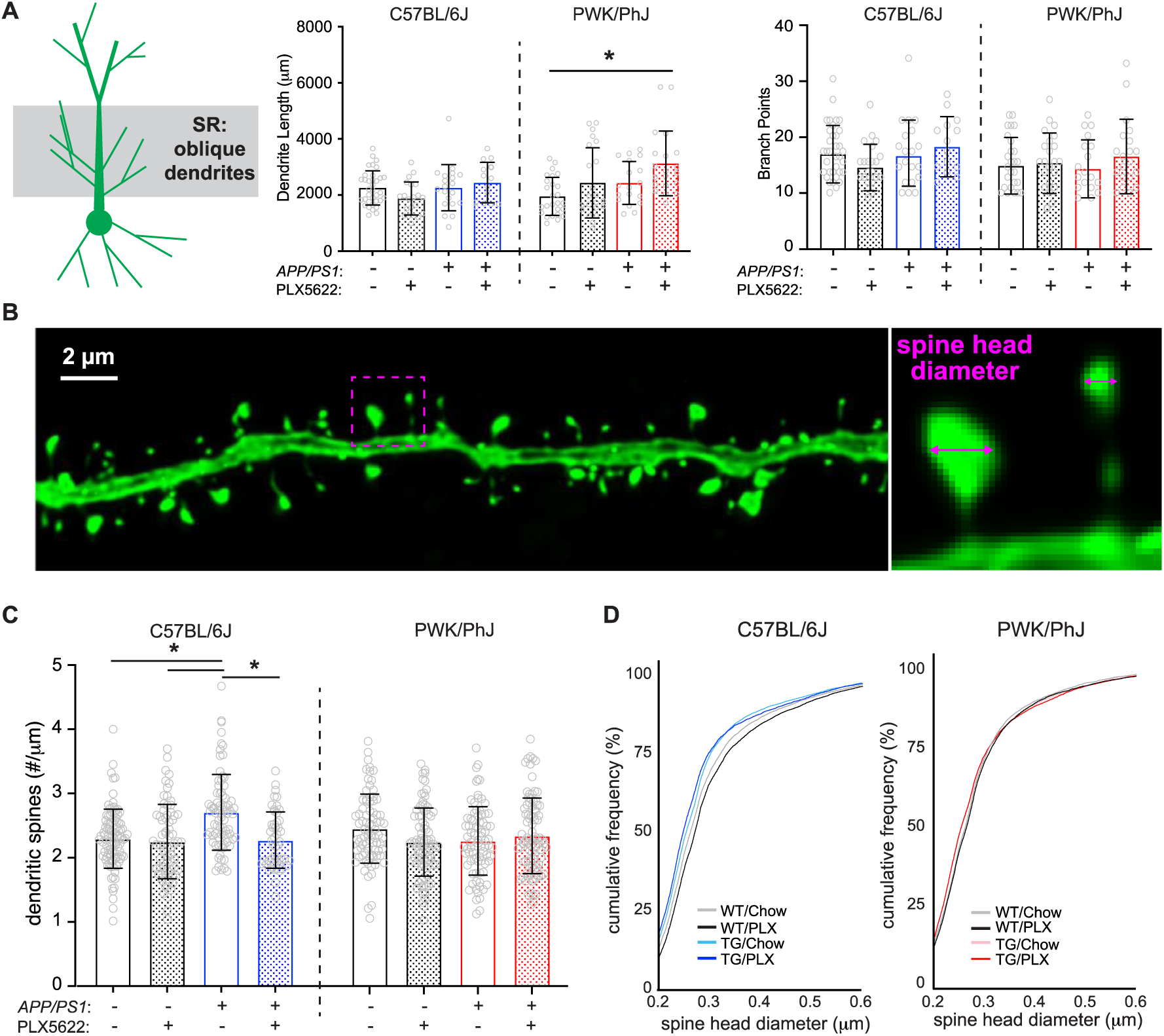
Amyloid- and microglia-dependent spine plasticity in oblique branches from B6 but not PWK mice. **(A)** Oblique dendritic lengths (left) and branch points (right). Individual data points represent each reconstructed neuron (n=3-5/mouse); error bars are ± SD; asterisks denote post-hoc (p<0.05) after two-way ANOVA and corrections for multiple comparisons. **(B)** Example deconvolved confocal image of an EGFP+ oblique branch. Spine density was acquired across dendrites (left) and individual spines measured for maximum head diameter (right). **(C)** Oblique branch spine densities across genotype/treatment groups. Data points represent individual branches (n=10-15/mouse); error bars are ± SD; asterisks denote comparisons (p<0.05) identified between control and PLX5622 groups after two-way ANOVA and corrections for multiple comparisons. **(D)** Spine head diameter cumulative distributions from B6 (left) and PWK (right). Kolmogorov-Smirnov (K-S) tests were used to evaluate statistical significance (see **Table S3**). Statistical analyses performed on B6 and PWK separately. For (A) and (C) *adjusted p<0.05 Bonferroni post-hoc tests (**Table S3**).

Synapse morphology is a reliable predictor of synaptic stability and strength^22, 23^ so we analyzed the maximum head diameter of each reconstructed spine. Comparison of spine sizes across the B6 groups revealed two prominent changes relative to the spine size distribution of WT control mice: a leftward shift in distribution of B6 TG control and TG PLX5622 compared to B6 WT control – indicating a population dominated by smaller spines, and a rightward shift in B6 WT PLX5622 compared to WT control – indicating a population dominated by larger spines. Like the spine density results, these shifts in spine sizes were noticeably absent across dendrites from PWK mice (**Figure 2D**). The opposing shifts in spine sizes across B6 mice was also evident in quartile-based density analyses (**Figure S2C**). The same quartile analysis from PWK mice revealed no differences, indicating a remarkable strain-specific stability of spine morphology to Aβ pathology or microglia depletion.

Basal branches in SO receive the same afferents from hippocampal area CA3 as the apical oblique branches in SR. The changes observed in apical oblique dendrites and dendritic spines were recapitulated in analyses of the basal dendrites in SO from these same projection neurons (**Figure S3**), including the relative maintenance of dendritic architecture, the TG-dependent increase in spine densities in B6 mice, and the morphological shift to smaller spine sizes between B6 WT control and TG control mice. Like the oblique dendrites, none of these effects were evident in basal dendrites from PWK mice. These results collectively show that B6 mice are vulnerable to Aβ-dependent changes in oblique (and to a lesser degree basal) spine density and morphology, while spines on dendrites from PWK mice appear resilient to Aβ pathology (**Figure S3, Table S4**).

### Differential patterns of spine loss or spine remodeling on the distal CA1 tuft dendrites

The distal dendrites of CA1 pyramidal cells receive synaptic input from neurons in the entorhinal cortex (EC), and these synapses show distinct morpho-molecular properties relative to synapses in SR and SO^35^. Although these inputs are more strongly filtered than those made onto the basal or oblique branches, they nevertheless are critical contributors to feature-selective firing of CA1 pyramidal cells^36, 37^. Like the maintenance of dendritic morphology in SR and SO, we found no significant effects on dendritic length across B6 and PWK groups except for a significant treatment effect on B6 branch points (**Table S5, Figure 3A**). Additional post-hoc and Sholl analyses suggested no two groups differed (**Figure S4A**).

**Figure 3:**
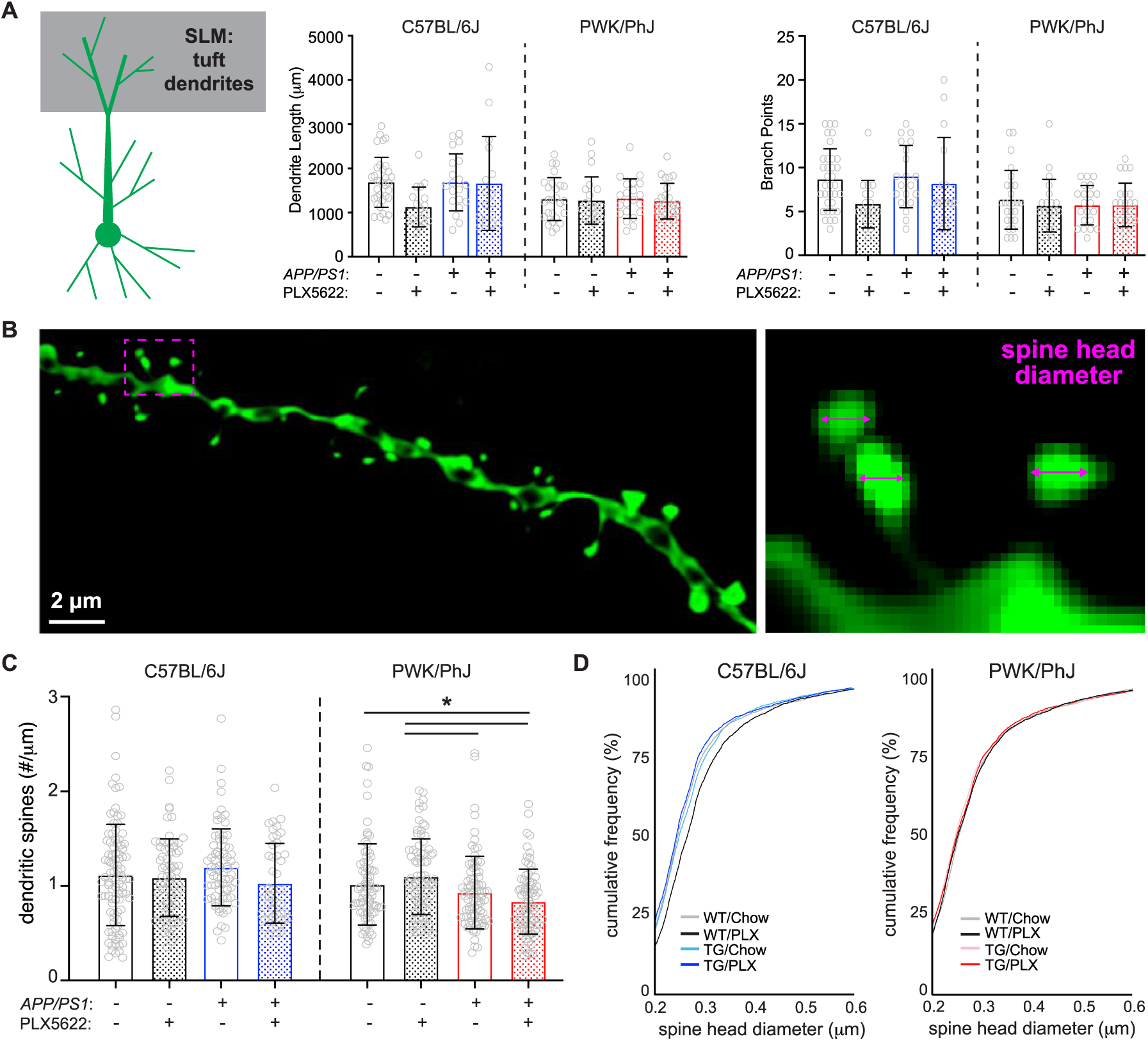
Differential regulation of spine density and morphology by amyloid on tuft branches from B6 and PWK *APP/PS1* mice. **(A)** Tuft dendritic lengths (left) and branch points (right). Individual data points represent each reconstructed neuron (n=3-5/mouse); error bars are ± SD; asterisks denote post-hoc (p<0.05) after two-way ANOVA and corrections for multiple comparisons. **(B)** Example deconvolved confocal image of an EGFP+ tuft branch segment. Spine density was acquired across dendrites (left) and individual spines measured for maximum head diameter (right). **(C)** Tuft branch spine densities across genotype/treatment groups. Data points represent individual branches (n=10-15/mouse); error bars are ± SD; asterisks denote comparisons (p<0.05) identified between PLX5622 and control diet groups after two-way ANOVA and corrections for multiple comparisons. **(D)** Spine head diameter cumulative distributions from B6 (left) and PWK (right). K-S tests were used to evaluate statistical differences (see **Table S5**). Statistical analyses performed on B6 and PWK separately. For (A) and (C) *adjusted p<0.05 Bonferroni post-hoc tests (**Table S5**).

Spines on tuft dendrites are lower in density but larger than those on basal or oblique (**Figure 3B**)^24^, suggesting these spines may be more stable as a population than those found on proximal SR/SO branches. Our measured tuft spine densities from B6 WT control mice recapitulated those obtained from AT and ssEM reconstructions from B6 mice (**Figure S4B**). In contrast to the effects observed in SR and SO, we observed no significant effects on spine densities from dendrites of B6 mice irrespective of genotype or treatment. In PWK mice, we found significant genotype and interactive effects, with TG control and TG PLX5622 mice exhibiting significantly lower spine densities than WT counterparts (**Table S5, Figure 3C**).

We next analyzed spine morphologies on these branches and observed a similar pattern of spine redistributions as those found on more proximal branches in B6 and PWK mice. Spines on branches from B6 TG control mice exhibited a leftward shift, indicating Aβ resulted in a population of smaller spines, while B6 WT PLX5622 branches shifted rightward, indicating microglia depletion resulted in a population of larger spines. Like the results from SR and SO, the experimental PWK groups showed no significant size redistributions (**Figure 3D, Table S5**), further supporting the resilience of this strain to Aβ or microglia activity. Quartile-based analysis further validated these population shifts in B6 but not PWK (**Figure S4C**). The disassociation between microglia-independent forms of amyloid-induced spine remodeling in B6, and spine loss in PWK, further supports the notion that these two strains are inherently different in their neuronal responses to Aβ pathology.

### Microglia-dendrite interactions influence spine density and size over large spatial scales

Microglia can influence neuronal synapses broadly through the release of diffusible messengers, or locally at points of microglia-dendrite ineractions^5, 6, 38^. To establish if the latter scenario was evident within our reconstructions, we examined spine data across dendritic segments that did (Touch+) or did not (Touch-) appear to physically interact with IBA1+ microglia (**Figure 4A**). Approximately 50% of dendrites from control diet mice were Touch-, whereas approximately 90% of dendrites from PLX5622 mice were Touch-(**Figure 4B, Table S6, Table S7**), providing a highly local estimate of depletion for our sampled branches. Given the sparsity of Touch+ dendrites in PLX5622 groups, analyses of microglia touch-based effects on spine density and size were only performed across control diet groups.

**Figure 4:**
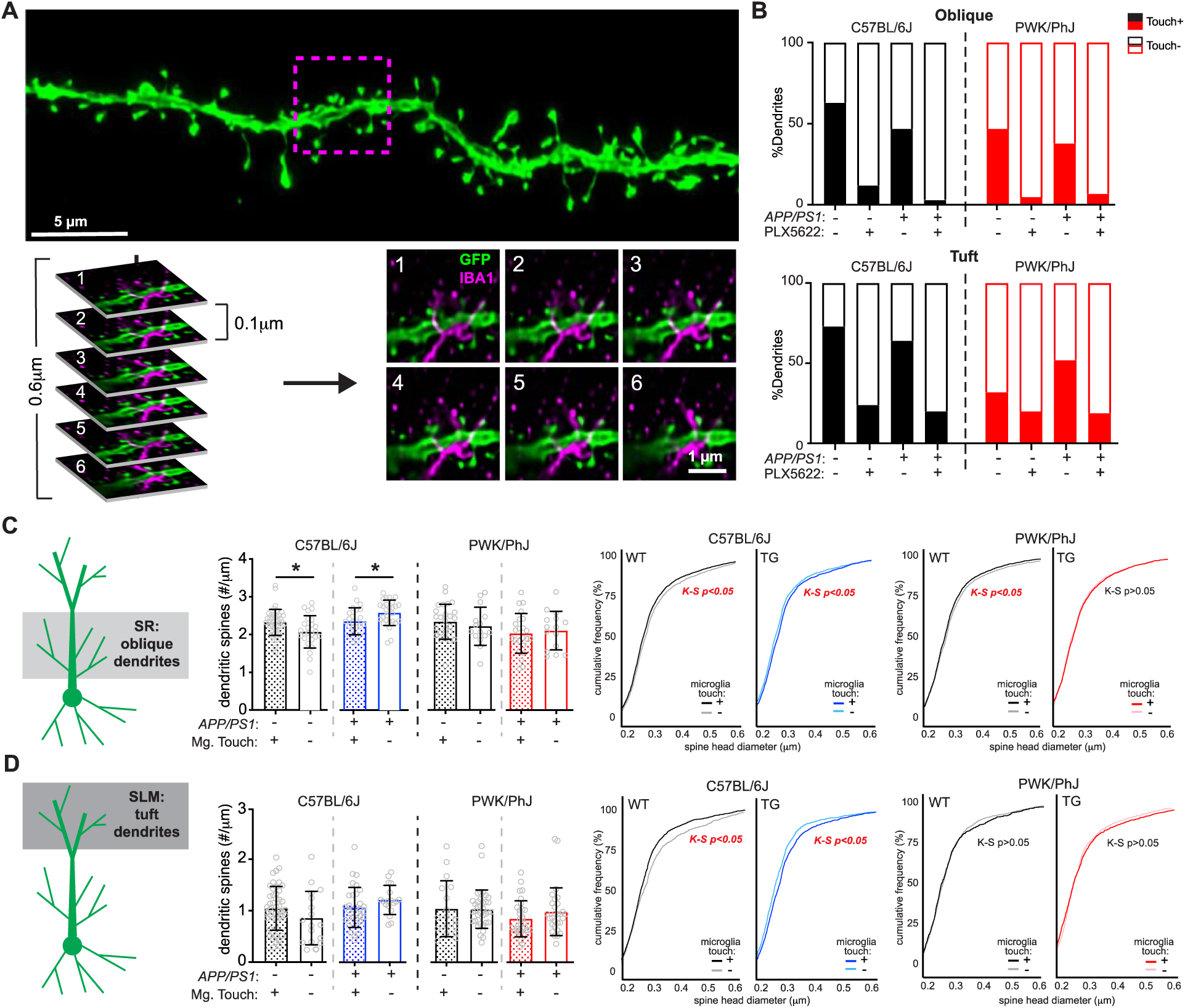
Microglia-dendrite interactions shape spine density and morphology. **(A)** Deconvolved confocal image of dendritic branch (green) with the region of microglia-dendrite interaction marked by a magenta box (top). Individual z-slices from the stack (0.1μm steps) with the IBA1+ microglia (in magenta) interacting with the EGFP+ dendrite (green) (bottom). Numbers in upper left hand denote z-steps. **(B)** Dendrite proportions that were classified as Touch+ or Touch-across oblique (top) and tuft (bottom) dendrites. **(C)** Spine densities (left) and cumulative distributions for spine head diameters (right) from oblique Touch+ and Touch-dendrites. Datapoints represent individual branches (n=5-10/mouse) and analyzed with unpaired t-tests between Touch+ and Touch-branches from B6 (middle) and PWK (right). K-S tests were used to evaluate statistical differences (see **Table S6**) **(D)** Identical analyses to (C) for tuft dendrites. See **Table S7** for statistics. For (C) and (D) *p<0.05, unpaired t-test.

In B6 WT mice, Touch+ oblique dendrites exhibited significantly higher spine densities compared to Touch-. Conversely, in B6 TG mice, Touch+ oblique dendrites showed significantly lower spine densities compared to Touch- (**Figure 4C**, left). These data suggest that in the absence of Aβ, microglia-dendrite interactions promote higher rates of synaptic connectivity, whereas microglia exposed to Aβ promote synaptic loss when contacting dendrites. Both effects were absent on dendrites from PWK mice, providing further evidence that genetic context controls how microglia regulate dendritic spines. Spines on Touch+ oblique dendrites from B6 WT mice were significantly smaller than those from Touch-, whereas spines on B6 TG Touch+ branches were significantly larger than Touch-. These patterns of microglia touch-dependent spine size changes were present on dendrites from PWK WT mice, but absent in PWK TG mice (**Figure 4C**, right). Thus, in terms of spine density and morphology, these data show that microglia-dendrite interactions regulate dendritic spines differently across B6 and PWK mice during Aβ plaque deposition.

Analysis of Touch+ and Touch-dendrites from the distal tuft compartment revealed no differences in dendritic spine density across B6 and PWK mice, regardless of genotype (**Figure 4D**, left). Like oblique branches, spines from B6 WT Touch+ tuft spines were significantly smaller than Touch-, while spines from B6 TG Touch+ were significantly larger than Touch-. Therefore, in B6 mice, microglia play opposing roles on spine morphology in healthy versus Aβ conditions. In contrast, PWK tuft spines exhibited no size differences between Touch+ and Touch-dendrites, regardless of genotype (**Figure 4D**, right).

Functional interactions among synapses can be highly localized within dendrites. For example, the induction of structural plasticity at one spine can lower the threshold for plasticity at neighboring spines within a restricted 5-10μm window^39, 40^. Therefore, we wondered if the effect of microglia Touch+ would be enhanced at the microglia-dendrite interaction point relative to adjacent locations on the same segment. Within Touch+ dendrites, we identified the point of microglia-dendrite interaction, and measured densities and head diameters of spines 5μm in each direction from the interaction point (proximal) and compared this to measures taken from adjacent 10μm-long dendritic section(s) (distal) (**Figure S5A**) from the same branch. Surprisingly, we did not identify significant differences in spine density and size between proximal and distal regions, regardless of strain or genotype (**Figure S5B**). The same analysis of distal tuft dendrites revealed a subtle yet significant increase in spine density in PWK WT animals at proximal compared to distal regions, but equivalent spine sizes across locations. Conversely, we found no differences in spine density at proximal versus distal regions in PWK TG mice but found that proximal spines were smaller than distal spines (**Figure S5C**). Thus, physical interactions between microglia and dendrites impact synaptic density and morphology over relatively large (mean segment length was 42 ± 4.9μm) spatial scales in B6 mice. Moreover, even when restricted to locations on branches where microglia physically interact with dendrites, responses in B6 mice still appear fundamentally different from PWK mice.

## DISCUSSION

We sought to determine how microglia regulate synapses on CA1 pyramidal neurons at an early phase of Aβ plaque deposition across genetically diverse contexts. Our analysis was restricted to CA1-to-PFC projecting pyramidal cells because they are the primary projection circuit to ventral PFC^41^, comprise an important pathway gating the progression of AD^42^, and allow for stringent comparisons within a single projection class across diverse mice. We took advantage of the same transgenic Aβ driver (*APP/PS1*) across genetically distinct B6 and PWK mouse strains and generating high-resolution reconstructions of ∼145,000 dendritic spine synapses. Since B6 and PWK mice exhibited equivalent rates of CA1 Aβ plaque deposition and microglia depletion, our results show that microglia-synapse interactions in healthy and Aβ-exposed brains depend on genetic context. Such a result strongly supports incorporating genetic diversity into mouse models of AD to faithfully recapitulate resilience, resistance, and susceptibility that are observed in the human patient population.

In B6 mice, CA1-to-PFC projection neurons showed significant changes in spine density and morphology on oblique branches in a microglia-dependent manner. More broadly, across all dendritic domains we found that *APP/PS1* and PLX5622 induced differential remodeling of spine sizes such that the overall spine population consisted of smaller (less stable^23, 24, 43^) synapses in the presence of Aβ plaque pathology, yet larger (more stable^23, 24, 43^) synapses in the absence of microglia. At a finer granularity, our results showed these changes in the B6 mice depend on whether physical interactions existed between microglial processes and individual pyramidal cell dendrites. The absence of each of these effects on CA1-to-PFC projection neuron dendrites from PWK mice provides the first evidence that like cognition^18^ and brain microglia^19^, synapses in PWK mice exhibit a form of resilience to Aβ plaque pathology. More broadly, these results strongly suggest that the mechanisms underlying synaptic changes in early AD vary across genetically diverse individuals. From a translational perspective, such a result suggests a low response rate to “one-size-fits-all” approaches to therapeutic interventions aimed at restoring synaptic structure and function during AD progression.

While our results agree with past work that reports that early synaptic changes in AD patients and mouse models may be subtle^29, 32, 44, 45^, more extreme rates of excitatory synapse loss in response to amyloid-driving AD transgenes have been reported^9, 33^. The differences between these studies and ours are likely due to several experimental factors (e.g., age at sacrifice, amyloid-driving transgene, methods estimating synaptic connections, and neurons examined). Being the first to examine Aβ- and microglia-dependent synaptic changes on specific projection neurons, it is possible that the CA1-to-PFC projection pathway we examined here shows a different pattern of synaptic responses than those on neurons chosen at random^9, 29^ or synapses sampled randomly from the neuronal population as a whole^32, 33^. Conversely, our rates of microglia depletion using PLX5622 were lower than those previously reported^10^. Yet we view this as a feature rather than a detriment, as this lowered depletion allowed us to measure microglia-dependent neuronal effects without influencing Aβ plaque pathology across *APP/PS1* mice.

One uniform feature across B6 and PWK mice was the heterogeneous laminar distribution of microglia within area CA1. The higher density of microglia in SLM was associated with elevated Aβ plaque burden which correlated with lower rates of PLX5622-mediated depletion, suggesting microglia in different hippocampal lamina may perform specific functions or belong to different transcriptionally-defined states^46^. Microglia states have gained increased interest with the initial discovery of disease-associated microglia (DAM) as the primary amyloid-induced state^47^ and interferon-responding microglia (IRM) as the primary aging-induced state^7^. Interestingly, we have found that B6.*APP/PS1* and PWK.*APP/PS1* female mice develop different proportions of these two microglia states^19^. Whether the differing susceptibility/resilience to CA1-to-PFC synaptic changes during Aβ plaque deposition across B6 and PWK *APP/PS1* mice seen here can be ascribed to specific microglia states should be determined through state-specific manipulation approaches.

The high-resolution but relatively low throughput assays used here, combined with the assessment of two genetically distinct mouse strains and two diet conditions, necessitated the use of only female mice in this study. Sex differences are important factors for dictating AD heterogeneity^48^, and have been reported in AD models including wild-derived mouse strains^18^. Similarly, we employed the *APP/PS1* transgene that is most relevant to Aβ deposition seen in familial AD^25, 49^, but the approach developed here can be applied more broadly to determine microglia-dependent synaptic changes across mouse models that are relevant to LOAD^50–52^. Despite these caveats, we show that the role of microglia in amyloid-induced synapse remodeling is highly dependent on genetic context. Intriguingly, PWK mice exhibit resilience that appears to stem from intrinsic neuronal mechanisms rather than protection from microglia. Identification of the genetic, cellular, and circuit-specific mechanisms of the resilience displayed in PWK mice could reveal novel therapeutic targets to prevent AD progression and promote early cognitive resilience across all patients. Alternatively, discovery of these neuronal resilience mechanisms could serve as a rational entry point towards precision therapeutic strategies in genetically defined subsets of AD patients.

## METHODS

## RESOURCE AVAILABILITY

### Lead Contact

Further information and requests for resources should be directed to and will be fulfilled by co-corresponding authors Erik Bloss, and Gareth Howell.

### Materials availability

All mouse strains are available through The Jackson Laboratory. All reagents in this study are commercially available.

### Data and code availability

Raw data and images from the figures will be available via Figshare (DOI provided upon publication).

## EXPERIMENTAL MODEL AND SUBJECT DETAILS

### Ethics Statement

All research was approved by the Institutional Animal Care and Use Committee (IACUC) at The Jackson Laboratory (approval number 12005 and 20006). Animals were humanely euthanized with 4% tribromoethanol (800mg/kg). Authors performed their work following guidelines established by “The Eight Edition of the Guide for the Care and Use of Laboratory Animals” and euthanasia using methods approved by the American Veterinary Medical Association.

### Mice

All mice were bred and housed in a 12/12 hour light/dark cycle on aspen bedding and fed a standard 6% Purina 5K52 Chow diet unless otherwise stated. Pilot experiments were performed on two mouse strains: C57BL/6J (JAX stock #000664) and PWK/PhJ (JAX stock #003715). Experimental cohorts were generated to produce 3 female mice per group. Mice were group housed for entirety of experiments. Primary experiments were performed on two additional mouse strains: B6.Cg-Tg(APPswe, PSEN1dE9)85Dbo/Mmjax (JAX stock #005864) and PWK.*APP/PS1* (JAX stock #25971). Experimental cohorts were generated to produce 6 female mice per group (12 *APP/PS1* carriers and littermate wild-type controls). However, due to increased seizure-induced mortality of B6.*APP/PS1* mice^53^, final cohorts for this strain were n=6 for TG control diet and n=5 for TG PLX5622 diet. Mice were initially group housed until 2.5 months of age (2 weeks before intracranial injections) mice were singly housed to avoid fighting-induced mortality among PWK.*APP/PS1* mice.

## METHOD DETAILS

### Intracranial viral injections

Recombinant viral vectors were used to drive Cre-recombinase (AAVretro-Cre), and Cre-dependent EGFP (serotype 2/1, AAV-flex-*rev*-EGFP)^20^. The titers of each virus were as follows (in genomic copies/mL): AAVretro-Cre, 1x10^12^; AAV-flex-*rev*-GFP, 1x10^13^. 30nL of AAVretro-Cre was injected into ventral prefrontal cortex (PFC) over 5 minutes, and 45-50nL (per each D/V coordinate) of AAV-flex-*rev*-GFP was injected in CA1 (CA1) over 10 minutes. Since PWK brain volumes are smaller than B6, injection coordinates were adjusted based on pilot experiments to determine injection sites. The coordinates for each injection were as follows (in mm: posterior relative to bregma, lateral relative to midline, and ventral relative to pial surface): B6 PFC (+1.75, -0.95, and -2.6), B6 CA1 (-3.5, -3.4, and -2.7/-2.5/2.0); PWK PFC (+1.45, -0.9, and -2.3), PWK CA1 (-3.5, -3.3, and -2.75/-2.5/-2.0). At each site the injection pipette was left in place for 3-5 minutes then slowly retracted at a rate of 10μm/s from the brain. After surgery mice were singly housed until sacrifice at 8 months of age.

### PLX5622 diet

PLX5622 was acquired from Chemgood (C-1521) and formulated in Purina 5K52 mouse chow diet at a concentration of 1200mg/kg (ppm) by Research Diets Inc, followed by 10-20 kGy gamma irradiation. Chemical purity and proper diet concentration were validated through HPLC and mass spectrometry analysis through Chemgood and JAX metabolomics core (detected 1030mg/kg purified from 1 pellet of diet). Mice were placed on diet at 4 months of age (4m) and left of diet until 8 months of age (8m). Mice were monitored weekly for food consumption and weighed monthly.

### Tissue harvest and brain sectioning

Mice were euthanized with an intraperitoneal lethal dose of Tribromoethanol (800mg/kg), followed by transcardial perfusion with 45mL ice-cold 4% paraformaldehyde (PFA) in 0.1M phosphate-buffered saline, in accordance with IACUC protocols. Brains were removed and placed in 5mL ice cold 4% PFA at 4°C for 24 hours, then placed into storage buffer (1XPBS + 0.1% Sodium Azide) for long-term storage at 4°C. Brains were sectioned at alternating thicknesses of 200μm and 50μm and kept in storage buffer (1XPBS + 0.1%NaN_3_) at 4°C until needed for imaging.

### Immunofluorescence analyses of microglia and amyloid-β plaques

50μm sections with EGFP+ dendrites were permeabilized with 1XPBS + 1% TritonX-100 (PBT), blocked for 12 hours at 4°C in PBT + 10% normal donkey/goat serum, washed once with PBT, and incubated in primary antibody solution containing rabbit anti-IBA1 (1:300, Wako) or chicken anti-IBA1 (1:500, Synaptic Systems). After primary incubation for 72 hours at 4°C, sections were washed 3 times with PBT and incubated in secondary antibodies (goat anti-rabbit Alexa Fluor 568 or donkey anti-chicken Alexa Fluor 647 (both diluted 1:500 in PBT)) for 24 hours at 4°C. Sections were washed with PBT, counterstained with DAPI (0.2mg/mL solution diluted 1:1000 in PBS), washed with PBS, and mounted with Vectashield Hardset mounting media. For assessing amyloid-β plaque pathology, additional 50μm sections underwent similar protocols, with X34 steps occurring before primary antibodies were applied. X34 solution was prepared by diluting 0.4mg X-34 (Sigma) in 4mL 200 proof ethanol, and 6 mL distilled water (diH_2_O). Sections were incubated in X-34 solution for 10 minutes, rinsed in diH_2_O for 3 minutes, incubated in 0.02M sodium hydroxide (NaOH) for 5 minutes, and washed in PBS. X34 staining was then proceeded by primary antibody staining. Additionally, since X34 and DAPI fluorescence are in overlapping channels, sections that underwent X34 staining were counterstained with TOPRO3 (1:1000 diluted in PBS) instead of DAPI.

Images were captured using two methodologies. EGFP+ sections that were stained with anti-IBA1 and DAPI were imaged on a Leica SP8 confocal microscope at 40X magnification, with each tile captured at 512 x 512 pixel frames using 2μm z-stack sizes. Sections that were stained for X34, anti-IBA1 and TOPRO3 were imaged on a Leica Versa slide scanner at 10X magnification, capturing and merging individual tiles. Analysis was performed using ImageJ2 (version 2.9.0/1.53t), where regions of interest (ROI) were outlined for SLM, SR, and SO as structurally visualized by the DAPI/TOPRO3 counterstains. For IBA1+DAPI+ quantification, maximum projections were generated from stacks, individual channels isolated, and default thresholds applied for IBA1 and DAPI to create binary images. IBA1 and DAPI binary images were merged for overlapping signal, followed by quantification of spots using the particle analyzer function (size threshold of 5-infinity pixels). For X34 quantification, the X34 channel was isolated from each image, thresholded to create a binary image, and quantified for total X34+ area and plaque number using the particle analyzer function. For quantification of plaque associated (PAM) and non-plaque associated (NPAM) microglia, 100μm in diameter ROIs were outlined around areas with X34+ plaques in SLM of each *APP/PS1* mouse. In the thresholded IBA1 channel from each imaged brain, IBA1+ area was quantified for each PAM ROI using the particle analyzer function. The sum of total IBA1+ area from each measured SLM was obtained and subtracted from the total quantified IBA1+ area to obtain NPAM IBA1+ area. When multiple SLM were analyzed per mouse, mean values from each PAM and NPAM ROI were calculated to obtain individual mouse statistics. Each quantified measure was normalized to area of ROI to obtain accurate densities.

### CAA Scoring

CAA severity was semi-quantitatively evaluated as described previously^26^. Images of X34+ plaques from transgenic *APP/PS1* mice were evaluated for CAA by three individual scorers, each blinded to the strain and treatment. Each image was assigned a semi-quantitative score ranging from 0 to 4 by the criteria as follows: 0 = no amyloid in vessels, 0.5 = scattered amyloid observed in leptomeninges, 1 = scattered amyloid in leptomeningeal and cortical vessels, 2 = strong circumferential amyloid deposition in multiple cortical and leptomeningeal vessels, 3 = widespread strong amyloid deposition in leptomeningeal and cortical vessels, and 4 = extravasation of amyloid deposition accompanied by dysphoric amyloid. For each image, the mode of the three scorers was obtained. If multiple images were acquired for each mouse, the mean CAA score was calculated to obtain a representative mouse score.

### Dendrite imaging, reconstruction, and analysis

200μm sections containing EGFP+ dendrites were identified using a fluorescence dissecting light microscope, were mounted on slides with 2 stacked 120μm imaging spacers (Electron Microscopy Sciences) and coverslipped in Vectashield mounting media. Images were acquired on a Leica SP8 confocal microscope at 40X magnification (oil immersion), 512 x 512 pixel dimensions, 1.25X digital zoom, 2μm z-steps, and depth-dependent detector gain compensation to maintain signal strength through the depth of the stack. Images were converted into TIFFs, followed by dendritic reconstructions in NeuronStudio^54^ (v0.9.92) software. For dendrite reconstructions, a maximum of 5 dendritic arbors from each compartment (e.g. basal, apical oblique, tuft) per mouse were reconstructed. Dendritic compartments were defined as follows: basal dendritic compartments included multiple origins, each of which emanate from the soma and traverse away from the SR; oblique dendritic compartments originate as a thick primary branch from the soma and gives rise to multiple terminal oblique dendrites; and tuft dendritic compartments originate at the site of the primary apical dendrite bifurcation as the dendrite enters the SLM and continues at inidivdual branch termination. After reconstructions, three dimensional Sholl analysis was performed on each dendrite using concentric circles spaced 20μm apart, originating at the soma for basal and oblique, and at the primary bifurcation for tuft dendrites. Summary statistics including total dendritic length and number of branch points were also acquired from this analysis, and each reconstructed neuron was reported as an individual measure.

### CA1-to-PFC dendritic spine imaging and analysis

EGFP+ dendrites were imaged on an SP8 confocal microscope equipped with a 63X objective (oil immersion), images collected at 50nm pixel sizes with 0.1μm z-steps, and stacks deconvolved using Leica Lightning software. 5-15 dendrites per compartment (e.g. basal, apical oblique, tuft) were captured per mouse. Slices that underwent co-labeling with anti-IBA1 antibodies had an additional channel captured for the secondary antibody signals. Each image was exported as TIFF format and imported into NeuronStudio^54^ for analysis of dendritic spine densities and morphologies. Density measurements were acquired by first reconstructing the dendritic cable followed by semi-automated spine identification. Cumulative distributions of assigned spine head diameters were analyzed by Kolmogorov-Smirnov tests, and through a quartile-based analysis. In this latter analysis, spines within dendritic compartments from each strain were pooled across treatment groups to create a population, and the first and last quartiles determined. From each branch, spines belonging to the first quartile (Q1, smallest) and last quartile (Q4, largest) were identified and density for each quartile per branch was calculated. Data were analyzed with each dendrite representing an individual data point.

### Microglia touch: proximal versus distal analysis

Images from 50μm sections that were co-labeled with anti-IBA1 were assessed for microglia-dendrite interactions by merging the EGFP (488) and IBA1 (568/647) channels in each slice in the z-stack. Images were then classified as Touch+ or Touch-based on whether the dendritic signal was physically overlapping with an IBA1+ process. Within the group of images that were Touch+, the exact x-y-z coordinates of the point of interaction were identified, and spine densities/morphologies calculated for the region of the dendrite 5μm on either side of the location of the touch (proximal, 10μm dendritic segment total). Spine densities and morphologies were gathered for the dendritic region that was 10μm adjacent to the dendritic region proximal to the microglia contact (distal). If the microglia contact appeared in the center of the dendrite, two distal 10μm dendritic regions were created on either side of the proximal zone, and mean spine density/morphology from distal zones calculated so that each proximal dendrite included a paired corresponding distal dendrite. Spine densities were calculated by each reconstructed dendrite, and spine sizes analyzed by each measured spine.

### Pilot study: tissue harvest and brain sectioning

Mice were euthanized with an intraperitoneal lethal dose of ketamine/xylazine (10 mg ketamine, 2 mg xylazine in 0.1mL sterile pure water per 10g body weight), followed by cardiac puncture and transcardial perfusion with 1XPBS in accordance with IACUC protocols. Blood collected from cardiac puncture was placed in EDTA-coated microtubes at room temperature until processing for flow cytometric analysis. Brains were removed and hemisected. Left brain hemispheres were placed in ice cold homogenization solution (Hank’s balanced salt solution [HBSS] with 15mM HEPES and 0.5% glucose), and immediately processed for flow cytometric analysis. Right brain hemispheres were placed in 5mL ice cold 4% paraformaldehyde (PFA) at 4°C for 24 hours, 10% sucrose for 24 hours, 30% sucrose for 24 hours, frozen and stored at -80°C for long-term storage.

### Pilot study: brain homogenization, myeloid cell preparation and FACs analysis

Brains were homogenized and myeloid cells isolated as described previously^19^. All hemispheres were homogenized on ice and performed using ice cold solutions to avoid myeloid cell activation. Each hemisphere was minced using a scalpel, followed by homogenization with a 15mL PTFE tissue grinder (4-5 strokes) in 2mL homogenization buffer. The suspension was transferred to a 50mL tube and passed through a pre-wet (with homogenization buffer) 70μm strainer. The suspension was transferred to a 15mL tube and spun in a centrifuge at 500g for 5 minutes at 4°C. After discarding the supernatant the cell pellet was resuspended in 2mL MACS buffer (PBS + 5% bovine serum albumin (BSA) + 2mM Ultrapure EDTA) for myelin removal procedure. 200μL Myelin Removal Beads (Miltenyi Biotec) was added to the cell suspension and mixed by gently pipetting. The cell suspension was divided equally into two 2mL microcentrifuge tubes and incubated at 4°C for 10 minutes. After incubation 1mL MACS buffer was added to each tube and centrifuged for 30s at 9300g at 4°C. After discarding supernatant, cell pellets were resuspended in 1.5mL MACS buffer per tube, and transferred to two pre-wet LD columns (Miltenyi Biotec). The flow through was collected in 50mL tubes on ice, and LD columns rinsed twice with 2mL MACS buffer. The final flow-through with washes were divided into multiple 2mL centrifuge tubes and centrifuged at 9300g for 30s at 4°C. After discarding supernatants, cell pellets were resuspended in 1mL 1XPBS. Since additional debris was still present, samples underwent debris removal protocol. After resuspension wit 1XPBS, samples were transferred to 15mL conical tubes and 900μL Debris Removal solution (Miltenyi Biotec) was added to each tube. Each mixture was carefully and slowly overlayed with 4mL 1XPBS, and centrifuged at 3000g for 4 minutes at 4°C. The top 2 interfaces were removed from the density gradient formed after centrifugation, and the bottom layer was saved on ice. 10mL 1XPBS was added to each tube, inverted gently, and centrifuged at 1000g for 10 minutes at 4°C. Supernatants were removed, and pellets resuspended in 1mL 1XPBS and stored on ice. Flow flow cytometric analysis, each sample was stained with DAPI, CD45 BV605 (clone 30-F11, BD Biosciences, 1:240), and CD11b PE (clone M1/70, Biolegend, 1:960), followed by processing on a FACSymphony A5 cytometer, and analysis using FlowJo (v10) software.

### Pilot study: blood preparation and FACs analysis

Blood collected from cardiac punctures were first processed by lysing red blood cells, followed by staining with the following antibodies: CD11c FITC (clone N418, TONBO, 1:600), CD19 PerCP-Cy5.5 (clone 1D3, TONBO, 1:480), CD11b PE (clone M1/70, Biolegend, 1:960), CD3e PE-CD594 (clone 145-2C11, BD Biosciences, 1:120), CD62L PE-Cy7 (clone MEL-14, TONBO, 1:600), CD4 APC (clone RM4-5, Biolegend, 1:480), CD8a A700 (clone 53-6.7, Biolegend, 1:600), Ly6G BV421 (clone 1A8, BD Biosciences, 1:480), CD45 BV605 (clone 30-F11, BD Biosciences, 1:240), B220 BUV496 (clone RA3-6B2, BD Biosciences, 1:120). Samples were processed on a FACSymphony A5 cytometer and analyzed using FlowJo (v10) software.

### Pilot study: immunofluorescence and image analysis

Brain hemispheres were sectioned at 25μm thickness and stored at 4°C in cryopreservative solution of 37.5% 1XPBS, 31.25% glycerol and 31.25% ethylene glycol. Sections were permeabilized with PBT, blocked for 12 hours at 4°C in PBT + 10% normal donkey serum, washed once with PBT, and incubated in primary antibody solution containing rabbit anti-IBA1 (1:300, Wako). After primary incubation for 48 hours at 4°C, sections were washed 3 times with PBT and incubated in secondary antibodies (donkey anti-rabbit Alexa Fluor 548868 (diluted 1:500 in PBT)) for 2 hours at room temperature. Sections were washed with PBT, counterstained with DAPI (0.2mg/mL solution diluted 1:1000 in PBS), washed with PBS, and mounted with Vectashield Hardset mounting media. Imaging was performed on a Leica SP8 confocal microscope at 20X magnification (glycerol immersion), 1024 x 1024 dimensions, using 1μm z-stack sizes. IMARIS software was used for analysis (v9.5.1) by creating a pseudo-channel with colocalized DAPI and IBA1 signal, and quantifying puncta using Spots function.

## QUANTIFICATION AND STATISTICAL ANALYSIS

### Statistical analysis

Data were analyzed blinded to genotype and treatment group. All statistical analyses were performed in GraphPad Prism software (v9.5.1) except for Kolmogorov-Smirnov tests which were performed using R (v4.2.2). Results are reported in table form in the Supplemental Information. Data from B6 and PWK mouse strains were analyzed separately. To assess treatment and genotype effects within each strain, two-way ANOVAs were computed followed by Bonferroni post hoc tests. Differences between treatment groups from plaque associated and non-plaque associated microglia area were assessed using nonparametric two-tailed t-tests within each strain. Comparisons of spine densities from this study to previously published array tomography and electron microscopy findings were tested using one-way ANOVA followed by Bonferroni post hoc tests. Quartile-based analyses to tests for differences in Q1 and Q4 densities between quartiles across groups were performed within strain with one-way ANOVA followed by Bonferroni post-hoc tests. Within-quartile effects on spine density across genotype/treatment groups were assessed with two-way ANOVA within strains followed by Bonferroni post-hoc tests. To determine differences in spine density within strain/treatment/genotype group based on microglia contact (Touch+ vs/ Touch-), two-tailed nonparametric unpaired t-tests were performed. Within a dendritic segment containing and microglial contact, spine densities between proximal versus distal regions in relation to microglial touch were evaluated using nonparametric two-tailed paired t-tests within strain/treatment/genotype group.

## ACKNOWLEDGEMENTS

We thank Will Schott, Danielle Littlefield, and Krystal-Leigh Brown in the Flow Cytometry core facility at The Jackson Laboratory for their expertise and assistance; Dr. Brian Hoffmann at The Jackson Laboratory for conducting metabolomic studies of PLX5622 diet; and Dr. Philipp Henrich at the Microscopy core at The Jackson Laboratory for training and assistance. We thank Melanie Maddocks Goodrich for assistance with producing and maintaining mouse experimental cohorts. This study was supported by the Diana Davis Spencer Foundation (G.R.H.), by the NIA AG079877 (E.B.B.), and by the NIA T32 training program AG062409 in the Precision Genetics of Aging, Alzheimer’s Disease and Related Dementias at The Jackson Laboratory (S.E.H., G.R.H., E.B.B.).

## AUTHOR CONTRIBUTIONS

S.E.H., E.B.B. and G.R.H. designed the study, with input from K.D.O. and K.C.G. A.A.H. and K.J.K. generated mouse cohorts and assisted with animal harvests and tissue collection. S.E.H. maintained mouse cohorts, performed all associated experiments, collected, and analyzed final data. E.B.B. provided training in intracranial injections, confocal microscopy imaging of dendrites and dendritic spines, and NeuronStudio analysis. E.B.B. and G.R.H. advised on all data analysis. K.D.O. and K.C.G. advised on data interpretation and manuscript preparation. S.E.H., E.B.B. and G.R.H. wrote the manuscript. All authors approved the final version.

