## Supplemental Information for "Genetic context controls early microglia-synaptic interactions in mouse models of Alzheimer’s disease"

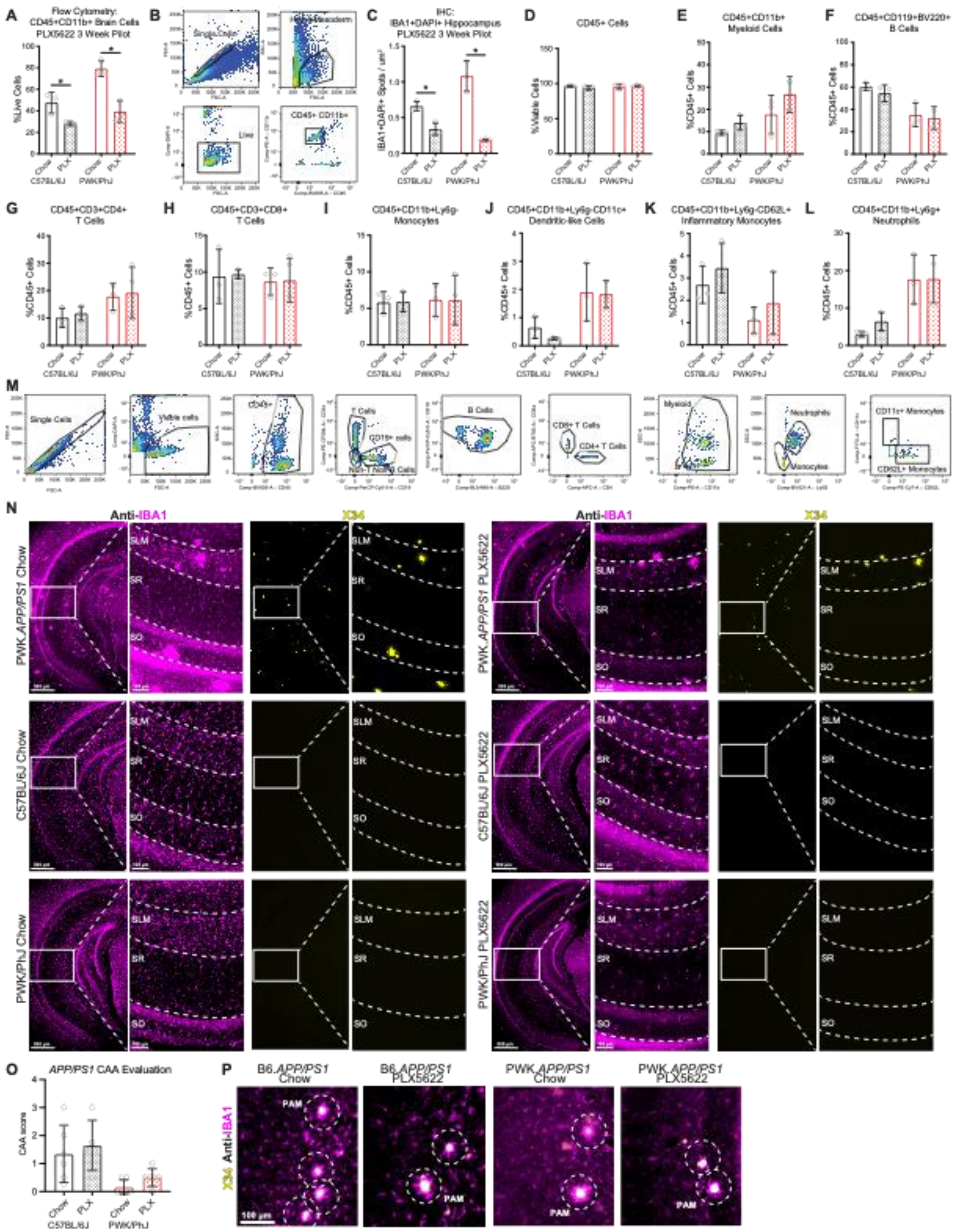
A
Flow Cytometry: CD45+CD11b+ Brain Cells PLX5622 3 Week Pilot
%Live Cells
Chow
PLX
C57BL/6J
PWK/PhJ
B
Single Cells
CD45+ CD11b+
Live
C
IHC: IBA1+DAPI+ Hippocampus PLX5622 3 Week Pilot
D
CD45+ Cells
%Viable Cells
E
CD45+CD11b+ Myeloid Cells
%CD45+ Cells
F
CD45+CD119+BV220+ B Cells
G
CD45+CD3+CD4+ T Cells
H
CD45+CD3+CD8+ T Cells
I
CD45+CD11b+Ly6g- Monocytes
J
CD45+CD11b+Ly6g-CD11c+ Dendritic-like Cells
K
CD45+CD11b+Ly6g-CD62L+ Inflammatory Monocytes
L
CD45+CD11b+Ly6g+ Neutrophils
M
Single Cells
Viable cells
cd45+
T Cells
CD19+ cells
B Cells
CD8+ T Cells
CD4+ T Cells
Myeloid
Neutrophils
Monocytes
CD11c+ Monocytes
CD62L+ Monocytes
N
Anti-IBA1
X34
PWK.APP/PS1 Chow
PWK.APP/PS1 PLX5622
C57BL/6J Chow
C57BL/6J PLX5622
PWK/PhJ Chow
PWK/PhJ PLX5622
SLM
SR
SO
O
APP/PS1 CAA Evaluation
CAA score
P
B6.APP/PS1 Chow
B6.APP/PS1 PLX5622
PWK.APP/PS1 Chow
PWK.APP/PS1 PLX5622
X34 Anti-IBA1
PAM
100 µm

**Figure S1: Evaluation of PLX5622-mediated microglia depletion across B6 and PWK mice, related to Figure 1**.

**(A-B)** Flow cytometric analysis (A) and gating strategy (B) of CD45+CD11b+ cells isolated from brain hemispheres of B6 and PWK mice treated for 3 weeks with chow diet or PLX5622 diet. Data presented as percent (%) of live cells. Two-way ANOVA detected significant ($p<0.05$) treatment, strain, and interactions.

**(C)** Immunohistochemical analysis of CA1 anti-IBA1 fluorescence from B6 and PWK mice treated for 3 weeks with chow control or PLX5622 diet. Two-way ANOVA detected significant ($p<0.05$) treatment and interactions.

**(D-M)** Flow cytometric analysis of peripheral blood from B6 and PWK female mice treated with chow diet or PLX5622 diet for 3 weeks. Blood cell populations were quantified: CD45+ cells (D), CD45+CD11b+ myeloid cells (E), CD45+CD19+BV220+ B cells (F), CD45+CD3+CD4+ helper T cells (G), CD45+CD3+CD8+ cytotoxic T cells (H), CD45+CD11b+Ly6g- monocytes (I), CD45+CD11v+Ly6g-CD11c+ dendritic-like cells (J), CD45+CD11b+Ly6g-CD62L+ inflammatory monocytes (K), and CD45+CD11b+Ly6g+ neutrophils. CD45+ cells in (D) reported as percent (%) of live cells. Populations quantified in (E-L) reported as % of CD45+ cells. Example gating strategy for peripheral blood analysis is depicted in (M). Two-way ANOVA identified significant ($p<0.05$) strain effects for (E), (F), (G), (J), (K), and (L).

**(N)** Images of IBA1+ microglia (magenta) and X34+ A$\beta$ plaques (yellow) across CA1 sub-regions part of the long-term PLX5622 study. SLM = stratum lacunosum moleculare, SR = stratum radiatum, SO = stratum oriens.

**(O)** Scoring of CAA severity across *APP/PS1* mice. Nonparametric two-tailed t-tests identified no significant within-strain differences between control chow and PLX5622 treated animals (see **Table S2**).

**(P)** Images corresponding to **Figure 1** analysis of plaque-associated microglia (PAM) across *APP/PS1* mice. Images taken at 10X magnification, and regions of interest (100$\mu$m in diameter) identified as X34+ and X34- across the SLM (5/mouse) in X34 channel. Separately, circular regions overlayed on corresponding IBA1 channels, and IBA1+ area quantified for each region. Depicted images are merged IBA1 (magenta) and X34 (yellow) 10X images, with example circular ROIs outlined for PAM.

All data presented at mean ± SD, with individual measures plotted as grey points.

Statistical analyses performed on B6 and PWK mouse strains together. For (A), (C)-(L) *adjusted $p<0.05$ Bonferroni post-hoc tests (see **Table S1**).

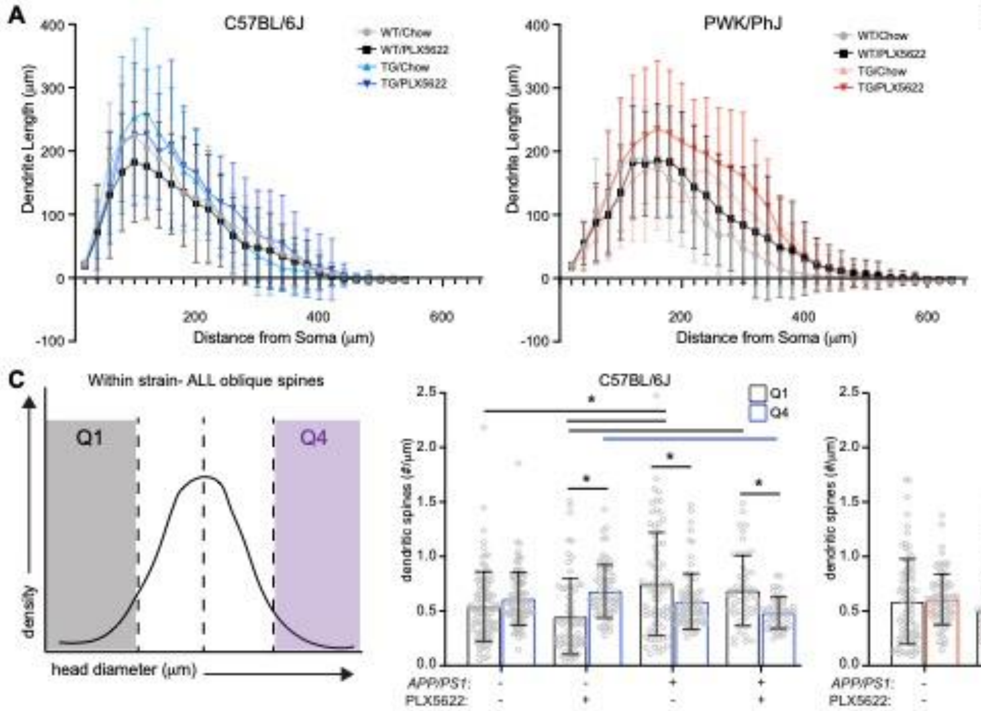


**Figure S2: Analyses of oblique dendrite structural organization and spine size analysis, related to Figure 2**.

**(A)** Sholl analysis of oblique dendrites reporting dendritic length (μm) in 20μm concentric distances from the soma origin. Data presented as mean ± SD of reconstructed neurons (n=3-5/mouse). Two-way ANOVA of dendrite length identify significant ($p<0.05$) genotype effect 120μm and 160-200μm away from the neuronal soma in B6. In PWK significant treatment identified at 220μm and 300-340μm, significant genotype 220-380μm, and significant interaction at 100μm, 380μm and 480-500μm from soma.

**(B)** Comparisons of oblique spine densities acquired from B6 WT/+ chow animals from the current study to array tomography (AT) and serial section electron microscopy (ssEM) oblique densities acquired from previous studies[28,35] with young, B6 mice. Data on graph is presented as mean ± SD, with data points representing measures from individual branches. One-way ANOVA identified significant ($p<0.05$) effect with $F = 6.578$.

**(C)** Quartile-based analyses of oblique spine head diameters. All oblique spines within each strain were divided into quartiles based on head diameter (μm). The smallest spines assigned to the first quartile (black, Q1) and the largest spines assigned to the fourth quartile (blue/red, Q4) were identified and reassigned back to originating dendrite. Spine densities (spines/μm) for Q1 and Q4 spines were calculated separately. Data points represent individual oblique branches. Two-way ANOVA within Q1 identified significant ($p<0.05$) genotype effect, and within Q4 identified significant genotype and interaction. One-way ANOVA across quartiles identified significant effect in B6 ($F=7.582$) with no effect in PWK.

All data in bar graphs presented at mean ± SD, with individual measures plotted as grey points.

Statistical analyses (unless noted otherwise) performed on B6 and PWK mouse strains separately. For (B) and (C) *adjusted $p<0.05$ Bonferroni post-hoc tests (see **Table S3**).

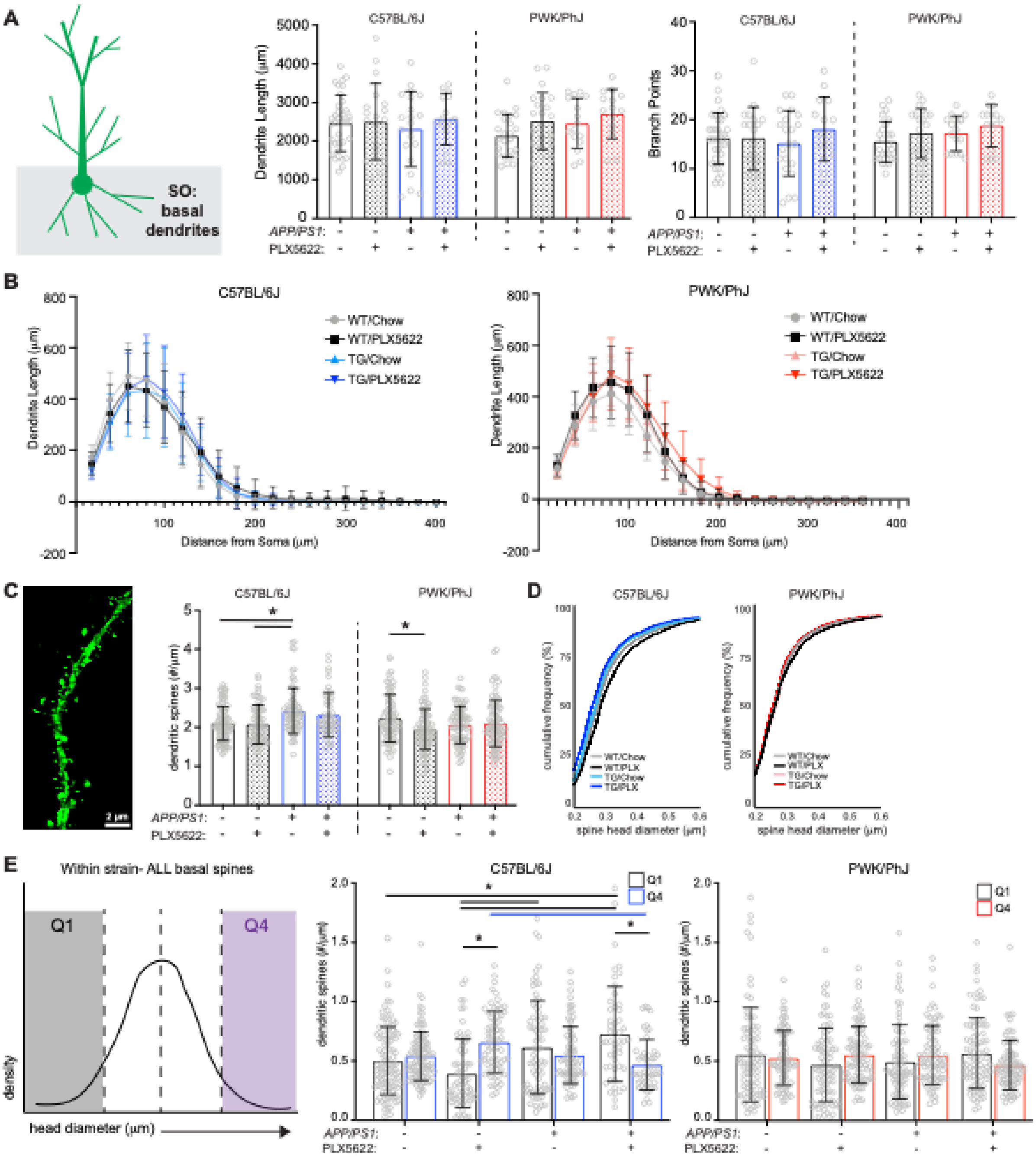


**Figure S3: Basal dendritic structure, spine density and spine size, related to Figure 2.**

**(A)** Basal dendritic reconstruction analyses quantifying total dendritic length (left) and number of branch points (right) across the dendritic tree. GFP+ dendrites were manually reconstructed from confocal images acquired at 40X magnification. Reconstructions were assigned an origin at the neuronal soma and traced all basal branches until point of termination. Individual data points represent each reconstructed neuron ($n \leq 5$/mouse). Two-way ANOVA for total dendrite length identify significant ($p<0.05$) treatment effects for PWK only.

**(B)** Sholl analysis of basal dendrites reporting dendritic length (μm) in 20μm concentric distances from the origin set at the neuronal soma. Data on graph is presented as mean ± SD of reconstructed neurons (n=3-5/mouse). Two-way ANOVA of dendrite length identify significant ($p<0.05$) treatment effect at 20μm, and genotype effect 20-40μm from neuronal soma in B6. In PWK significant treatment at 120-200μm, genotype at 140-240μm, and interaction at 60μm and 180μm from soma.

**(C)** Image of a basal EGFP+ branch (left), and results of basal spine densities (spines/μm; center) across genotype/treatment groups. Data points represent individual branches ($n \leq 15$/mouse) (right). Two-way ANOVA identify significant ($p<0.05$) genotype in B6, and significant treatment effect and interaction in PWK.

**(D)** Cumulative distributions of spine head diameters (μm) for basal spines across genotype/treatment groups, separated by strain. Statistical results from Kolmogorov-Smirnov (K-S) tests for differences in cumulative distributions are reported in **Table S4**.

**(E)** Quartile-based analysis of basal spine head diameters (performed as in **S2C**). Two-way ANOVA in B6 identified significant ($p<0.05$) genotype effects and interaction in both Q1 and Q4, and in PWK identified significant interactions in both Q1 and Q4. One-way ANOVA across quartiles identified significant effect in B6 ($F=8.564$) with no effect in PWK.

All data in bar graphs presented at mean ± SD, with individual measures plotted as grey points.

Statistical analyses (unless noted otherwise) performed on B6 and PWK mouse strains separately. For (A), (C), (E) *adjusted $p<0.05$ Bonferroni post-hoc tests (see **Table S4**).

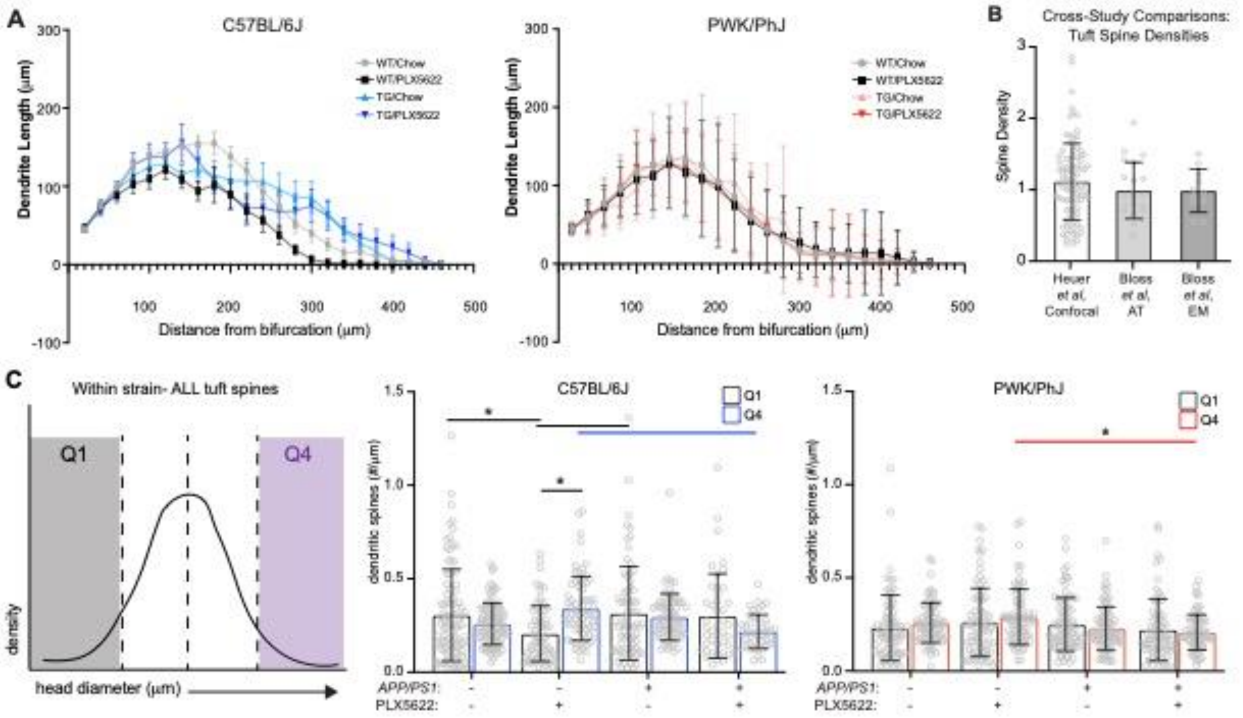


**Figure S4: Analyses of tuft dendritic structure and spine sizes, related to Figure 3**.

**(A)** Sholl analysis of tuft dendrites reporting dendritic length (µm) in 20µm concentric distances from the origin set at the main bifurcation of the primary apical branch. Data on graph is presented as mean ± SD of reconstructed neurons (n=3-5/mouse). Two-way ANOVA of dendrite length identify significant ($p<0.05$) genotype effect 220µm, treatment effect 280-380µm, and interaction 140-160µm from main bifurcation in B6 only.

**(B)** Cross-study comparisons of tuft spine densities acquired from B6 WT/+ chow animals from the current study (using confocal microscopy), AT and ssEM tuft densities acquired from young, B6 mice[28,35]. Data on graph is presented as mean ± SD, with data points representing measures from individual branches. One-way ANOVA identified no significant effect.

**(C)** Quartile-based analysis of tuft spine head diameters (as in **S2C**). Two-way ANOVA within Q1 identified significant ($p<0.05$) treatment effect, and within Q4 identified significant genotype and interaction in B6, and significant genotype effect within Q4 in PWK. One-way ANOVA across quartiles identified significant effects in both B6 ($F=4.419$) and PWK ($F=2.666$).

All data in bar graphs presented at mean ± SD, with individual measures plotted as grey points. Statistical analyses (unless noted otherwise) performed on B6 and PWK mouse strains separately. For (C) *adjusted $p<0.05$ Bonferroni post-hoc tests (see **Table S5**).

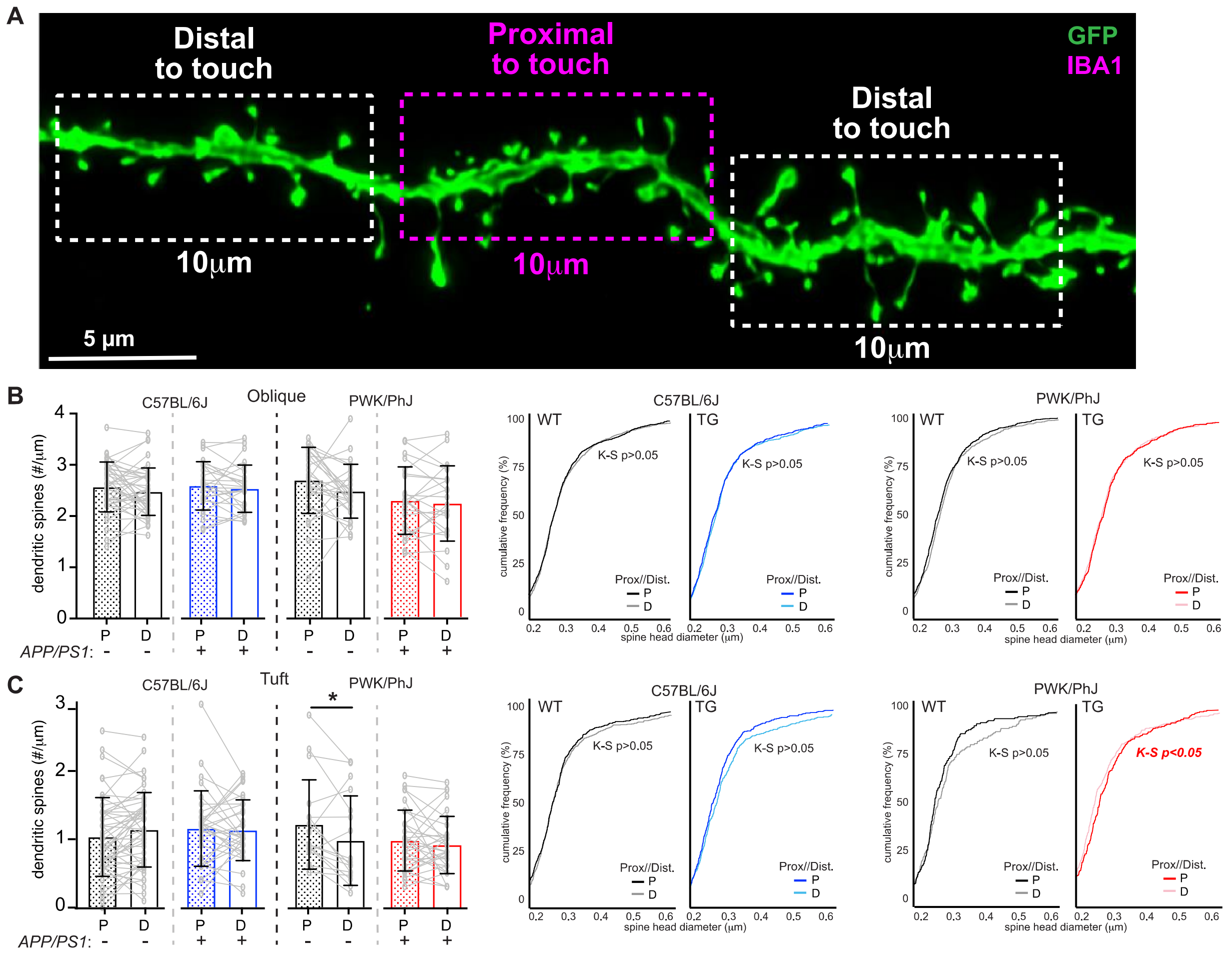


**Figure S5: Microglia-dendrite interactions do not selectively shape spines at the point of contact, related to Figure 4**.

**(A)** Image of the dendrite depicted in **Figure 4A**, with outlines depicting regions that are proximal and distal to a microglia-dendrite interaction (Touch+). 10μm of each type of segment for each dendrite with a microglia contact were quantified for spine density and head diameter.

**(B)** Analysis of spine density (left) and head diameter (right) of Touch+ oblique branches, comparing region of the dendrite proximal versus distal to the microglia contact. Spine density data points represent individual branches (n≤15 mouse). Paired t-tests were performed to compare spine densities between proximal and distal branch segments within each strain/genotype group.

**(C)** Identical analysis to (B) for tuft dendrites.

For (B) and (C) *$p<0.05$, paired t-test. Cumulative distributions were statistically analyzed with Kolmogorov-Smirnov (K-S) tests. All statistical analyses for corresponding diagrams reported in **Table S6** and **Table S7**.

### SUPPLEMENTAL TABLES

### Table S1- Associated Statistics for 3 Week PLX Pilot, related to Figure S1

#### S1A. CD45+CD11b+ Brain Myeloid Cells Flow Cytometry

| Strain | Chow vs. PLX5622 Bonferroni post-hoc t-test adj. p-value |
|---|---|
| C57BL/6J | 0.0342 |
| PWK/PhJ | 0.0005 |
| Two-Way ANOVA p-value: | Treatment: 0.0002<br>Strain: 0.0017<br>Interaction: 0.0560 |

#### S1B. IBA1+DAPI+ / um$^2$ CA1 IHC

| Strain | Chow vs. PLX5622 Bonferroni post-hoc t-test adj. p-value |
|---|---|
| C57BL/6J | 0.0215 |
| PWK/PhJ | <0.0001 |
| Two-Way ANOVA p-value: | Treatment: <0.0001<br>Strain: 0.0892<br>Interaction: 0.0031 |

#### S1C. %Microglia Depletion- Flow and IHC

| | % Depletion ((PLX-chow)/chow)*100 | |
|---|---|---|
| **Strain** | **Flow<br>Brain Hemisphere<br>%live CD45+CD11b+ cells** | **IHC<br>Dorsal CA1<br>IBA1+DAPI+ cells** |
| B6 | 40.8 ± 7.4 | 49.0 ± 8.1 |
| PWK | 50.4 ± 7.8 | 83.2 ± 2.1 |

#### S1D. Peripheral Blood Flow Cytometry

| | Chow vs. PLX5622 Bonferroni post-hoc t-test adj. p-value | | |
|---|---|---|---|
| **Blood Cell Population (%CD45+ Cells)** | **C57BL/6J** | **PWK/PhJ** | **Two-Way ANOVA p-values** |
| CD45+ Cells (%Viable) | 0.55570 | >0.9999 | Treatment: 0.6083<br>Strain: 0.5238<br>Interaction: 0.2989 |
| CD45+CD11b+<br>Myeloid | 0.8018 | 0.2229 | Treatment: 0.0951<br>Strain: 0.0185<br>Interaction: 0.5415 |
| CD45+CD19+BV220+<br>B Cells | 0.8323 | >0.9999 | Treatment: 0.4001<br>Strain: 0.001<br>Interaction: 0.7544 |
| CD45+CD3+CD4+<br>T Cells | >0.9999 | >0.9999 | Treatment: 0.6633<br>Strain: 0.0497<br>Interaction: 0.9793 |
| CD45+CD3+CD8+<br>T Cells | >0.9999 | >0.9999 | Treatment: 0.8820<br>Strain: 0.6184<br>Interaction: 0.9599 |
| CD45+CD11b+Ly6g-<br>Monocytes | >0.9999 | >0.9999 | Treatment: 0.9822<br>Strain: 0.8318<br>Interaction: 0.9625 |
| CD45+CD11b+ly6g-CD11c+<br>Dendritic-like Cells | 0.9129 | >0.9999 | Treatment: 0.5371<br>Strain: 0.0035<br>Interaction: 0.6564 |
| CD45+CD11b+Ly6g-CD62L+<br>Inflammatory Monocytes | 0.8043 | 0.7764 | Treatment: 0.2395<br>Strain: 0.0292<br>Interaction: 0.9848 |
| CD45+CD11b+Ly6g+<br>Neutrophils | 0.8128 | >0.9999 | Treatment: 0.5376<br>Strain: 0.0015<br>Interaction: 0.5681 |

### S1E. Body Weight

| | Chow vs. PLX5622<br>Bonferroni post-hoc t-test adj. p-value | | |
|---|---|---|---|
| **Treatment Week** | **C57BL/6J** | **PWK/PhJ** | **Two-Way ANOVA p-values** |
| Week 0 | >0.9999 | 0.9241 | Treatment: 0.4577<br>Strain: <0.0001<br>Interaction: 0.7629 |
| Week 1 | 0.8826 | >0.9999 | Treatment: 0.5272<br>Strain: <0.0001<br>Interaction: 0.6409 |
| Week 2 | >0.9999 | 0.8918 | Treatment: 0.3790<br>Strain: 0.0002<br>Interaction: 0.8446 |
| Week 3 | 0.1408 | 0.0960 | Treatment: 0.0141<br>Strain: <0.0001<br>Interaction: 0.8665 |

### S1F. Food Consumption

| | Chow vs. PLX5622<br>Bonferroni post-hoc adj. p-value | | |
|---|---|---|---|
| **Treatment Week** | **C57BL/6J** | **PWK/PhJ** | **Two-Way ANOVA p-values** |
| Week 1 | >0.9999 | >0.9999 | Treatment: 0.2487<br>Strain: 0.2217 |
| Week 2 | >0.9999 | >0.9999 | Treatment: 0.6625<br>Strain: 0.1498 |
| Week 3 | >0.9999 | >0.9999 | Treatment: 0.5806<br>Strain: 0.2557 |

### Table S2- Associated Statistics for IBA1+DAPI+ and X34 Counts, related to Figure 1 and Figure S1

#### S2A. B6 IBA1+DAPI+ / um$^2$ ANOVA followed by Bonferroni post-hoc pairwise analysis

| C57BL/6J | | SLM | SR | SO |
|---|---|---|---|---|
| Group A | Group B | Adj. p-value | Adj. p-value | Adj. p-value |
| WT/+ Chow | WT/+ PLX5622 | <0.0001 | <0.0001 | <0.0001 |
| WT/+ Chow | TG/+ Chow | 0.0419 | >0.9999 | >0.9999 |
| WT/+ Chow | TG/+ PLX5622 | <0.0001 | <0.0001 | <0.0001 |
| WT/+ PLX5622 | TG/+ Chow | <0.0001 | <0.0001 | <0.0001 |
| WT/+ PLX5622 | TG/+ PLX5622 | 0.8012 | >0.9999 | >0.9999 |
| TG/+ Chow | TG/+ PLX5622 | <0.0001 | <0.0001 | <0.0001 |
| Two-way ANOVA within vCA1 region (p-values): | | Treatment: <0.0001 Genotype: 0.0209 Interaction: 0.5283 | Treatment: <0.0001 Genotype: 0.9898 Interaction: 0.7809 | Treatment: <0.0001 Genotype: 0.07532 Interaction: 0.05250 |
| Cross-region two-way ANOVA (p-values): | | vCA1 region: <0.0001 Genotype/Treatment group: <0.0001 Interaction: <0.0001 | | |

#### S2B. PWK IBA1+DAPI+ / um$^2$ ANOVA followed by Bonferroni post-hoc pairwise analysis

| PWK/PhJ | | SLM | SR | SO |
|---|---|---|---|---|
| Group A | Group B | Adj. p-value | Adj. p-value | Adj. p-value |
| WT/+ Chow | WT/+ PLX5622 | 0.0002 | 0.0005 | 0.0023 |
| WT/+ Chow | TG/+ Chow | <0.0001 | 0.0449 | 0.0119 |
| WT/+ Chow | TG/+ PLX5622 | >0.9999 | 0.0272 | 0.0378 |
| WT/+ PLX5622 | TG/+ Chow | <0.0001 | <0.0001 | <0.0001 |
| WT/+ PLX5622 | TG/+ PLX5622 | 0.0125 | >0.9999 | >0.9999 |
| TG/+ Chow | TG/+ PLX5622 | <0.0001 | <0.0001 | <0.0001 |
| Two-way ANOVA within vCA1 region (p-values): | | Treatment: <0.0001 Genotype: <0.0001 Interaction: 0.1562 | Treatment: <0.0001 Genotype: 0.0008 Interaction: 0.1674 | Treatment: <0.0001 Genotype: 0.7979 Interaction: 0.8307 |
| Cross-region two-way ANOVA (p-values): | | vCA1 region: <0.0001 Genotype/Treatment group: <0.0001 Interaction: 0.2072 | | |

#### S2C. Microglia depletion efficiencies across vCA1 (% of chow control counterpart)

| | % Depletion ((PLX-chow)/chow)*100 | | |
|---|---|---|---|
| Strain/Genotype | SLM | SR | SO |
| B6 WT/+ | 61.4 ± 5.8 | 62.8 ± 3.9 | 60.9 ± 5.3 |
| B6 TG/+ | 58.9 ± 3.1 | 64.4 ± 3.5 | 59.9 ± 5.9 |
| PWK WT/+ | 48.8 ± 7.2 | 75.2 ± 6.5 | 61.1 ± 10.1 |
| PWK TG/+ | 46.7 ± 6.9 | 68.1 ± 7.4 | 64.6 ± 7.0 |

#### S2D. B6 X34 / um$^2$ ANOVA followed by Bonferroni post-hoc test

| C57BL/6J | | SLM | | SR | | SO | |
|---|---|---|---|---|---|---|---|
| | | Adj. p-value | | Adj. p-value | | Adj. p-value | |
| Group A | Group B | Area | Spots | Area | Spots | Area | Spots |
| TG/+ Chow | TG/+ PLX5622 | 0.4584 | >0.9999 | >0.9999 | >0.9999 | >0.9999 | 0.4550 |
| Two-Way ANOVA across vCA1 regions (p-values) | | **Area** vCA1 Region: <0.0001 Treatment: 0.2381 Interaction: 0.6295 | | | **Spots** vCA1 Region: 0.0146 Treatment: 0.7769 Interaction: 0.2838 | | |

### S2E. PWK X34 / um$^2$ ANOVA followed by Bonferroni post-hoc test

| PWK/PhJ | | SLM | | SR | | SO | |
|---|---|---|---|---|---|---|---|
| | | Adj. p-value | | Adj. p-value | | Adj. p-value | |
| **Group A** | **Group B** | **Area** | **Spots** | **Area** | **Spots** | **Area** | **Spots** |
| TG/+ Chow | TG/+ PLX5622 | >0.9999 | 0.1118 | >0.9999 | >0.9999 | >0.9999 | >0.9999 |
| Two-Way ANOVA across vCA1 regions (p-values) | | **Area**<br>vCA1 Region: <0.0001<br>Treatment: 0.9515<br>Interaction: 0.7969 | | | **Spots**<br>vCA1 Region: <0.0001<br>Treatment: 0.0907<br>Interaction: 0.3688 | | |

### S2F. Plaque-associated IBA1+ microglia area / um$^2$ nonparametric two-tailed t-test

| SLM | | B6.*APP/PS1* p-value | | PWK.*APP/PS1* p-value | |
|---|---|---|---|---|---|
| **Group A** | **Group B** | **PAM** | **NPAM** | **PAM** | **NPAM** |
| Chow | PLX5622 | 0.0303 | 0.0173 | 0.0022 | 0.0022 |

### S2G. Plaque-associated IBA1+ microglia %reduction between chow and PLX5622 groups

| SLM | | B6.*APP/PS1* %reduction | | PWK.*APP/PS1* %reduction | |
|---|---|---|---|---|---|
| **Group A** | **Group B** | **PAM** | **NPAM** | **PAM** | **NPAM** |
| Chow | PLX5622 | 43.196 ± 11.000 | 53.216 ± 9.205 | 48.856 ± 3.033 | 66.388 ± 7.832 |

### S2H. CAA Score statistics across *APP/PS1* mice

| Strain | Chow vs. PLX5622 Nonparametric t-test, p-value |
|---|---|
| C57BL/6J (TG) | 0.5173 |
| PWK/PhJ (TG) | 0.1775 |

### Table S3- Associated Statistics for Oblique Dendrites & Spine Densities, related to Figure 2 and S2

#### S3A. B6 Dendrite ANOVA with Bonferroni post-hoc tests

| C57BL/6J | | Length | Branches |
|---|---|---|---|
| **Group A** | **Group B** | **Adj. p-value** | **Adj. p-value** |
| WT/+ Chow | WT/+ PLX5622 | 0.2027 | 0.5157 |
| WT/+ Chow | TG/+ Chow | >0.9999 | >0.9999 |
| WT/+ Chow | TG/+ PLX5622 | >0.9999 | >0.9999 |
| WT/+ PLX5622 | TG/+ Chow | 0.2292 | >0.9999 |
| WT/+ PLX5622 | TG/+ PLX5622 | 0.0124 | 0.0587 |
| TG/+ Chow | TG/+ PLX5622 | >0.9999 | 0.9119 |
| Two-way ANOVA (p-values): | | Treatment: 0.5187<br>Genotype: 0.0595<br>Interaction: 0.0674 | Treatment: 0.7450<br>Genotype: 0.1511<br>Interaction: 0.0885 |

#### S3B. PWK Dendrite ANOVA with Bonferroni post-hoc tests

| PWK/PhJ | | Length | Branches |
|---|---|---|---|
| **Group A** | **Group B** | **Adj. p-value** | **Adj. p-value** |
| WT/+ Chow | WT/+ PLX5622 | 0.2705 | >0.9999 |
| WT/+ Chow | TG/+ Chow | 0.1902 | >0.9999 |
| WT/+ Chow | TG/+ PLX5622 | 0.0008 | 0.7707 |
| WT/+ PLX5622 | TG/+ Chow | >0.9999 | >0.9999 |
| WT/+ PLX5622 | TG/+ PLX5622 | 0.2311 | >0.9999 |
| TG/+ Chow | TG/+ PLX5622 | 0.5378 | >0.9999 |
| Two-way ANOVA (p-values): | | Treatment: 0.0073<br>Genotype: 0.0076<br>Interaction: 0.6187 | Treatment: 0.2693<br>Genotype: 0.7940<br>Interaction: 0.4755 |

#### S3C. Dendrite Sholl Analysis- ANOVA

| | Dendrite length: Two-way ANOVA | | | | | |
|---|---|---|---|---|---|---|
| | C57BL/6J | | | PWK/PhJ | | |
| **Distance from Soma (um)** | **Treatment p-value** | **Genotype p-value** | **Interaction p-value** | **Treatment p-value** | **Genotype p-value** | **Interaction p-value** |
| 20 | 0.3456 | 0.3120 | 0.5651 | 0.8721 | 0.9152 | 0.6875 |
| 40 | 0.3219 | 0.1775 | 0.3319 | 0.1582 | 0.2316 | 0.4282 |
| 60 | 0.0531 | 0.3126 | 0.3475 | 0.8697 | 0.0889 | 0.1597 |
| 80 | 0.2128 | 0.2247 | 0.4634 | 0.9184 | 0.7417 | 0.1087 |
| ***100*** | 0.1609 | 0.1354 | 0.7559 | 0.7745 | 0.8581 | 0.0196 |
| ***120*** | 0.1627 | 0.0206 | 0.9197 | 0.2812 | 0.8742 | 0.1881 |
| 140 | 0.1614 | 0.0502 | 0.9051 | 0.2757 | 0.5355 | 0.1353 |
| ***160*** | 0.7261 | 0.0316 | 0.4177 | 0.0925 | 0.2613 | 0.1962 |
| ***180*** | 0.7880 | 0.0448 | 0.8301 | 0.0623 | 0.0948 | 0.6075 |
| ***200*** | 0.9352 | 0.0287 | 0.5504 | 0.1507 | 0.0668 | 0.6441 |
| ***220*** | 0.9831 | 0.3360 | 0.6409 | 0.0375 | 0.0011 | 0.9724 |
| ***240*** | 0.6179 | 0.3423 | 0.3160 | 0.0594 | <0.0001 | 0.5475 |
| ***260*** | 0.4599 | 0.1797 | 0.0576 | 0.0534 | <0.0001 | 0.6876 |
| ***280*** | 0.4914 | 0.5131 | 0.0289 | 0.0707 | <0.0001 | 0.6897 |
| ***300*** | 0.7015 | 0.5893 | 0.0588 | 0.0144 | <0.0001 | 0.7756 |
| ***320*** | 0.4877 | 0.5533 | 0.0761 | 0.0318 | 0.0005 | 0.9032 |
| ***340*** | 0.1274 | 0.9745 | 0.0628 | 0.0200 | 0.0009 | 0.8915 |
| ***360*** | 0.4618 | 0.7213 | 0.1308 | 0.2116 | 0.0007 | 0.1505 |
| ***380*** | 0.3599 | 0.8126 | 0.5485 | 0.4468 | 0.0084 | 0.0224 |
| 400 | 0.6704 | 0.8113 | 0.3342 | 0.2260 | 0.0786 | 0.1599 |
| 420 | 0.6122 | 0.3768 | 0.2728 | 0.2654 | 0.1559 | 0.2570 |
| 440 | 0.9248 | 0.3012 | 0.2804 | 0.3052 | 0.5369 | 0.1961 |
| 460 | 0.6785 | 0.7177 | 0.1980 | 0.2755 | 0.8535 | 0.2831 |
| 480 | 0.9641 | 0.2899 | 0.3110 | 0.3222 | 0.5885 | 0.0896 |
| ***500*** | 0.1968 | 0.8605 | 0.8605 | 0.3711 | 0.7670 | 0.0427 |
| ***520*** | 0.2660 | 0.5813 | 0.5813 | 0.6999 | 0.6784 | 0.0373 |
| 540 | 0.5470 | 0.5470 | 0.5470 | 0.5453 | 0.8895 | 0.1056 |
| 560 | NA | NA | NA | 0.4470 | 0.4470 | 0.0828 |
| 580 | NA | NA | NA | 0.5691 | 0.5691 | 0.0773 |
| 600 | NA | NA | NA | 0.3793 | 0.3793 | 0.2981 |
| 620 | NA | NA | NA | 0.3360 | 0.3360 | 0.3360 |
| 640 | NA | NA | NA | 0.3360 | 0.3360 | 0.3360 |

### S3D. B6 Dendrite Sholl Analysis- Bonferroni post-hoc tests

| C57BL/6J | Bonferroni multiple comparison, adjusted p-value | | | | | |
|---|---|---|---|---|---|---|
| Distance from Soma (um) | WT/+ Chow vs. WT/+ PLX5622 | WT/+ Chow vs. TG/+ Chow | WT/+ Chow vs. TG/+ PLX5622 | WT/+ PLX5622 vs. TG/+ Chow | WT/+ PLX5622 vs. TG/+ PLX5622 | TG/+ Chow vs. TG/+ PLX5622 |
| 120 | 0.5535 | 0.0109 | >0.9999 | 0.0001 | 0.1200 | 0.7413 |
| 160 | 0.9781 | 0.6374 | 0.3329 | 0.0441 | 0.0247 | >0.9999 |
| 180 | >0.9999 | 0.3796 | 0.2227 | 0.6359 | 0.3727 | >0.9999 |
| 200 | >0.9999 | 0.6993 | 0.2906 | 0.3804 | 0.1607 | >0.9999 |

### S3E. PWK Dendrite Sholl Analysis- Bonferroni post-hoc tests

| PWK/PhJ | Bonferroni multiple comparison, adjusted p-value | | | | | |
|---|---|---|---|---|---|---|
| Distance from Soma (um) | WT/+ Chow vs. WT/+ PLX5622 | WT/+ Chow vs. TG/+ Chow | WT/+ Chow vs. TG/+ PLX5622 | WT/+ PLX5622 vs. TG/+ Chow | WT/+ PLX5622 vs. TG/+ PLX5622 | TG/+ Chow vs. TG/+ PLX5622 |
| 100 | 0.1117 | 0.0394 | >0.9999 | >0.9999 | 0.1220 | 0.0450 |
| 220 | 0.1691 | 0.0059 | <0.0001 | >0.9999 | 0.0082 | 0.3494 |
| 240 | 0.0648 | <0.0001 | <0.0001 | 0.2129 | 0.0040 | >0.9999 |
| 260 | 0.1307 | <0.0001 | <0.0001 | 0.1173 | 0.0008 | >0.9999 |
| 280 | 0.9478 | 0.0033 | <0.0001 | 0.2429 | 0.0002 | 0.2973 |
| 300 | 0.2062 | 0.0029 | <0.0001 | 0.8976 | 0.0004 | 0.1056 |
| 320 | 0.3070 | 0.0102 | <0.0001 | >0.9999 | 0.0052 | 0.3110 |
| 340 | 0.1961 | 0.0240 | <0.0001 | >0.9999 | 0.0501 | 0.5383 |
| 360 | 0.3727 | 0.0077 | 0.0085 | 0.9534 | >0.9999 | >0.9999 |
| 380 | 0.2957 | 0.0152 | 0.1912 | >0.9999 | >0.9999 | >0.9999 |
| 500 | >0.9999 | >0.9999 | >0.9999 | >0.9999 | >0.9999 | >0.9999 |
| 520 | >0.9999 | >0.9999 | >0.9999 | >0.9999 | >0.9999 | >0.9999 |

### S3F. B6 Spine Density & Head Diameter- ANOVA with Bonferroni post-hoc tests

| C57BL/6J | | Density | Head Diameter |
|---|---|---|---|
| Group A | Group B | Bonferroni post-hoc adj. p-value | Kolgorov-Smirnov p-value |
| WT/+ Chow | WT/+ PLX5622 | >0.9999 | 2.63e-11 |
| WT/+ Chow | TG/+ Chow | <0.0001 | 1.958e-13 |
| WT/+ Chow | TG/+ PLX5622 | >0.9999 | <2.2e-16 |
| WT/+ PLX5622 | TG/+ Chow | <0.0001 | <2.2e-16 |
| WT/+ PLX5622 | TG/+ PLX5622 | >0.9999 | <2.2e-16 |
| TG/+ Chow | TG/+ PLX5622 | <0.0001 | 4.135e-5 |
| Two-way ANOVA (p-values): | | Treatment: 0.0002<br>Genotype: 0.0006<br>Interaction: 0.0021 | |

### S3G. PWK Spine Density & Head Diameter- ANOVA with Bonferroni post-hoc tests

| PWK/PhJ | | Density | Head Diameter |
|---|---|---|---|
| Group A | Group B | Bonferroni post-hoc adj. p-value | Kolgorov-Smirnov p-value |
| WT/+ Chow | WT/+ PLX5622 | 0.0860 | 0.04846 |
| WT/+ Chow | TG/+ Chow | 0.1646 | 1.796e-7 |
| WT/+ Chow | TG/+ PLX5622 | >0.9999 | 6.651e-9 |
| WT/+ PLX5622 | TG/+ Chow | >0.9999 | 3.287e-11 |
| WT/+ PLX5622 | TG/+ PLX5622 | >0.9999 | 7.355e-13 |
| TG/+ Chow | TG/+ PLX5622 | >0.9999 | 0.6598 |
| Two-way ANOVA (p-values): | | Treatment: 0.2766<br>Genotype: 0.4333<br>Interaction: 0.0164 | |

### S3H. Comparison to AT and ssEM datasets

| Heuer vs. Bloss Comparisons | | Density |
|---|---|---|
| **Group A** | **Group B** | **Bonferroni post-hoc adj. p-value** |
| Heuer B6 WT/+ Chow Confocal | Bloss AT | 0.1797 |
| Heuer B6 WT/+ Chow Confocal | Bloss EM | 0.0034 |
| Bloss AT | Bloss EM | 0.1003 |
| One-Way ANOVA: | | F = 6.578<br>p-value = 0.0019 |

### S3I. B6 Spine head diameter within Quartile- ANOVA with Bonferroni post-hoc tests

| C57BL/6J | | Quartile | |
|---|---|---|---|
| **Group A** | **Group B** | **Q1<br>Bonferroni adj. p-value** | **Q4<br>Bonferroni adj. p-value** |
| WT/+ Chow | WT/+ PLX5622 | >0.9999 | >0.9999 |
| WT/+ Chow | TG/+ Chow | 0.0004 | >0.9999 |
| WT/+ Chow | TG/+ PLX5622 | 0.2039 | 0.4994 |
| WT/+ PLX5622 | TG/+ Chow | <0.0001 | >0.9999 |
| WT/+ PLX5622 | TG/+ PLX5622 | 0.0013 | 0.0219 |
| TG/+ Chow | TG/+ PLX5622 | >0.9999 | >0.9999 |
| Two-way ANOVA (p-value): | | Treatment: 0.0885<br>Genotype: <0.0001<br>Interaction: 0.7323 | Treatment: 0.5162<br>Genotype: 0.0001<br>Interaction: 0.0027 |

### S3J. PWK Spine head diameter within Quartile- ANOVA with Bonferroni post-hoc tests

| PWK/PhJ | | Quartile | |
|---|---|---|---|
| **Group A** | **Group B** | **Q1<br>Bonferroni adj. p-value** | **Q4<br>Bonferroni adj. p-value** |
| WT/+ Chow | WT/+ PLX5622 | >0.9999 | >0.9999 |
| WT/+ Chow | TG/+ Chow | >0.9999 | >0.9999 |
| WT/+ Chow | TG/+ PLX5622 | >0.9999 | >0.9999 |
| WT/+ PLX5622 | TG/+ Chow | >0.9999 | >0.9999 |
| WT/+ PLX5622 | TG/+ PLX5622 | 0.2133 | >0.9999 |
| TG/+ Chow | TG/+ PLX5622 | >0.9999 | >0.9999 |
| Two-way ANOVA (p-value): | | Treatment: 0.4410<br>Genotype: 0.0924<br>Interaction: 0.1469 | Treatment: 0.9811<br>Genotype: 0.1695<br>Interaction: 0.5977 |

### S3K. B6 between Quartiles- ANOVA with Bonferroni post-hoc test

| C57BL/6J | |
|---|---|
| **Group A** | **Q1 vs. Q4<br>Bonferroni adj. p-value** |
| WT/+ Chow | >0.9999 |
| WT/+ PLX5622 | 0.0004 |
| TG/+ Chow | 0.0319 |
| TG/+ PLX5622 | 0.0364 |
| One-way ANOVA<br>Quartile effect | F = 7.582<br>p-value = <0.0001 |

### S3L. PWK between Quartiles- ANOVA with Bonferroni post-hoc test

| PWK/PhJ | |
|---|---|
| **Group A** | **Q1 vs. Q4<br>Bonferroni adj. p-value** |
| WT/+ Chow | >0.9999 |
| WT/+ PLX5622 | >0.9999 |
| TG/+ Chow | >0.9999 |
| TG/+ PLX5622 | >0.9999 |
| One-way ANOVA<br>Quartile effect | F = 1.327<br>p-value = 0.2347 |

### Table S4- Associated Statistics for Basal Dendrites & Spine Densities, related to Figure S3

#### S4A. B6 Dendrites- ANOVA with Bonferroni post-hoc tests

| C57BL/6J | | Length | Branches |
|---|---|---|---|
| **Group A** | **Group B** | **Adj. p-value** | **Adj. p-value** |
| WT/+ Chow | WT/+ PLX5622 | >0.9999 | >0.9999 |
| WT/+ Chow | TG/+ Chow | >0.9999 | >0.9999 |
| WT/+ Chow | TG/+ PLX5622 | 0.8467 | 0.3642 |
| WT/+ PLX5622 | TG/+ Chow | >0.9999 | >0.9999 |
| WT/+ PLX5622 | TG/+ PLX5622 | >0.9999 | >0.9999 |
| TG/+ Chow | TG/+ PLX5622 | 0.8184 | 0.4025 |
| Two-way ANOVA (p-values): | | Treatment: 0.4257<br>Genotype: 0.8157<br>Interaction: 0.5642 | Treatment: 0.2716<br>Genotype: 0.7275<br>Interaction: 0.2858 |

#### S4B. PWK Dendrites- ANOVA with Bonferroni post-hoc tests

| PWK/PhJ | | Length | Branches |
|---|---|---|---|
| **Group A** | **Group B** | **Adj. p-value** | **Adj. p-value** |
| WT/+ Chow | WT/+ PLX5622 | 0.2575 | 0.6534 |
| WT/+ Chow | TG/+ Chow | >0.9999 | >0.9999 |
| WT/+ Chow | TG/+ PLX5622 | 0.2617 | 0.3339 |
| WT/+ PLX5622 | TG/+ Chow | >0.9999 | >0.9999 |
| WT/+ PLX5622 | TG/+ PLX5622 | >0.9999 | >0.9999 |
| TG/+ Chow | TG/+ PLX5622 | >0.9999 | >0.9999 |
| Two-way ANOVA (p-values): | | Treatment: 0.0425<br>Genotype: 0.0955<br>Interaction: 0.6250 | Treatment: 0.0966<br>Genotype: 0.1045<br>Interaction: 0.9313 |

#### S4C. Sholl Analysis- ANOVA

| | Dendrite length: Two-way ANOVA | | | | | |
|---|---|---|---|---|---|---|
| | C57BL/6J | | | PWK/PhJ | | |
| **Distance from Soma (um)** | **Treatment p-value** | **Genotype p-value** | **Interaction p-value** | **Treatment p-value** | **Genotype p-value** | **Interaction p-value** |
| ***20*** | 0.0129 | 0.0027 | 0.8502 | 0.8127 | 0.0732 | 0.2580 |
| ***40*** | 0.1682 | 0.0064 | 0.3409 | 0.5290 | 0.0908 | 0.0981 |
| 60 | 0.7205 | 0.2308 | 0.3894 | 0.8439 | 0.5658 | 0.0451 |
| 80 | 0.9588 | 0.9369 | 0.2163 | 0.2425 | 0.1813 | 0.7775 |
| 100 | 0.9674 | 0.2872 | 0.3741 | 0.1046 | 0.1441 | 0.4862 |
| ***120*** | 0.4708 | 0.1755 | 0.8365 | 0.0181 | 0.1437 | 0.6312 |
| ***140*** | 0.1560 | 0.2957 | 0.5433 | 0.0339 | 0.0473 | 0.6161 |
| ***160*** | 0.1113 | 0.7140 | 0.4928 | 0.0379 | 0.0238 | 0.1203 |
| ***180*** | 0.1127 | 0.6127 | 0.1813 | 0.0124 | 0.0065 | 0.0431 |
| ***200*** | 0.3624 | 0.5410 | 0.0903 | 0.0437 | 0.0047 | 0.4834 |
| ***220*** | 0.1513 | 0.4361 | 0.5428 | 0.8197 | 0.0232 | 0.3688 |
| ***240*** | 0.5390 | 0.4921 | 0.3200 | 0.8427 | 0.0320 | 0.7142 |
| 260 | 0.5395 | 0.2520 | 0.1899 | 0.7740 | 0.1615 | 0.5358 |
| 280 | 0.4096 | 0.1646 | 0.1450 | 0.9262 | 0.3359 | 0.4183 |
| 300 | 0.2670 | 0.2670 | 0.1220 | 0.3904 | 0.3904 | 0.3904 |
| 320 | 0.2633 | 0.2633 | 0.1461 | 0.3904 | 0.3904 | 0.3904 |
| 340 | 0.2294 | 0.2294 | 0.1964 | 0.3904 | 0.3904 | 0.3904 |
| 360 | 0.2577 | 0.2577 | 0.2577 | 0.3904 | 0.3904 | 0.3904 |
| 380 | 0.2577 | 0.2577 | 0.2577 | NA | NA | NA |
| 400 | 0.2577 | 0.2577 | 0.2577 | NA | NA | NA |

#### S4D. B6 Sholl Analysis- Bonferroni post-hoc tests

| C57BL/6J | Bonferroni multiple comparison, adjusted p-value | | | | | |
|---|---|---|---|---|---|---|
| **Distance from Soma (um)** | **WT/+ Chow vs. WT/+ PLX5622** | **WT/+ Chow vs. TG/+ Chow** | **WT/+ Chow vs. TG/+ PLX5622** | **WT/+ PLX5622 vs. TG/+ Chow** | **WT/+ PLX5622 vs. TG/+ PLX5622** | **TG/+ Chow vs. TG/+ PLX5622** |
| 20 | >0.9999 | >0.9999 | 0.2000 | >0.9999 | >0.9999 | >0.9999 |
| 40 | 0.1060 | 0.0007 | 0.0004 | >0.9999 | 0.7128 | >0.9999 |

### S4E. PWK Sholl Analysis- Bonferroni post-hoc tests

| PWK/PhJ | Tukey's multiple comparison, adjusted p-value | | | | | |
|---|---|---|---|---|---|---|
| **Distance from Soma (um)** | **WT/+ Chow vs. WT/+ PLX5622** | **WT/+ Chow vs. TG/+ Chow** | **WT/+ Chow vs. TG/+ PLX5622** | **WT/+ PLX5622 vs. TG/+ Chow** | **WT/+ PLX5622 vs. TG/+ PLX5622** | **TG/+ Chow vs. TG/+ PLX5622** |
| 60 | 0.0432 | 0.0246 | >0.9999 | >0.9999 | 0.7490 | 0.4303 |
| 120 | 0.0002 | 0.0664 | <0.0001 | >0.9999 | >0.9999 | 0.1518 |
| 140 | 0.2987 | 0.6113 | <0.0001 | >0.9999 | 0.0555 | 0.0622 |
| 160 | >0.9999 | >0.9999 | 0.0048 | >0.9999 | 0.0244 | 0.0658 |
| 180 | >0.9999 | >0.9999 | 0.0537 | >0.9999 | 0.1254 | 0.2434 |
| 200 | >0.9999 | >0.9999 | 0.6536 | >0.9999 | >0.9999 | >0.9999 |
| 220 | >0.9999 | >0.9999 | >0.9999 | >0.9999 | >0.9999 | >0.9999 |
| 240 | >0.9999 | >0.9999 | >0.9999 | >0.9999 | >0.9999 | >0.9999 |

### S4F. B6 Spine Density & Head Diameter- ANOVA with Bonferroni post-hoc tests

| C57BL/6J | | Density | Head Diameter |
|---|---|---|---|
| **Group A** | **Group B** | **Bonferroni post-hoc adj. p-value** | **Kolgorov-Smirnov p-value** |
| WT/+ Chow | WT/+ PLX5622 | >0.9999 | <2.2e-16 |
| WT/+ Chow | TG/+ Chow | 0.0003 | 1.265e-8 |
| WT/+ Chow | TG/+ PLX5622 | 0.0919 | <2.2e-16 |
| WT/+ PLX5622 | TG/+ Chow | 0.0005 | <2.2e-16 |
| WT/+ PLX5622 | TG/+ PLX5622 | 0.0852 | <2.2e-16 |
| TG/+ Chow | TG/+ PLX5622 | >0.9999 | 6.78e-13 |
| Two-way ANOVA (p-values): | | Treatment: 0.3085<br>Genotype: <0.0001<br>Interaction: 0.4816 | |

### S4G. PWK Spine Density & Head Diameter- ANOVA with Bonferroni post-hoc tests

| PWK/PhJ | | Density | Head Diameter |
|---|---|---|---|
| **Group A** | **Group B** | **Bonferroni post-hoc adj. p-value** | **Kolgorov-Smirnov p-value** |
| WT/+ Chow | WT/+ PLX5622 | 0.0082 | 3.821e-6 |
| WT/+ Chow | TG/+ Chow | 0.3521 | 8.563e-5 |
| WT/+ Chow | TG/+ PLX5622 | 0.7876 | 5.252e-5 |
| WT/+ PLX5622 | TG/+ Chow | >0.9999 | 0.1038 |
| WT/+ PLX5622 | TG/+ PLX5622 | 0.4759 | 1.675e-12 |
| TG/+ Chow | TG/+ PLX5622 | >0.9999 | 1.03e-11 |
| Two-way ANOVA (p-values): | | Treatment: 0.0445<br>Genotype: 0.8945<br>Interaction: 0.0101 | |

### S4H. B6 Spine Head Diameter within Quartile- ANOVA with Bonferroni post-hoc tests

| C57BL/6J | | Quartile (adj. p-value) | |
|---|---|---|---|
| **Group A** | **Group B** | **Q1** | **Q4** |
| WT/+ Chow | WT/+ PLX5622 | 0.5102 | 0.2366 |
| WT/+ Chow | TG/+ Chow | 0.2624 | >0.9999 |
| WT/+ Chow | TG/+ PLX5622 | 0.0002 | >0.9999 |
| WT/+ PLX5622 | TG/+ Chow | 0.0002 | 0.5898 |
| WT/+ PLX5622 | TG/+ PLX5622 | <0.0001 | 0.0100 |
| TG/+ Chow | TG/+ PLX5622 | 0.8933 | >0.9999 |
| Two-way ANOVA (p-value): | | Treatment: 0.9268<br>Genotype: <0.0001<br>Interaction: 0.0070 | Treatment: 0.5053<br>Genotype: 0.0009<br>Interaction: 0.0004 |

### S4I. PWK Spine Head Diameter within Quartile- ANOVA with Bonferroni post-hoc tests

| **PWK/PhJ** | | **Quartile (adj. p-value)** | |
|---|---|---|---|
| **Group A** | **Group B** | **Q1** | **Q4** |
| WT/+ Chow | WT/+ PLX5622 | >0.9999 | >0.9999 |
| WT/+ Chow | TG/+ Chow | >0.9999 | >0.9999 |
| WT/+ Chow | TG/+ PLX5622 | >0.9999 | >0.9999 |
| WT/+ PLX5622 | TG/+ Chow | >0.9999 | >0.9999 |
| WT/+ PLX5622 | TG/+ PLX5622 | 0.7335 | >0.9999 |
| TG/+ Chow | TG/+ PLX5622 | >0.9999 | >0.9999 |
| Two-way ANOVA (p-value): | | Treatment: 0.8729<br>Genotype: 0.6014<br>Interaction: 0.0313 | Treatment: 0.2608<br>Genotype: 0.2125<br>Interaction: 0.0353 |

### S4J. B6 between Quartiles- ANOVA with Bonferroni post-hoc tests

| **C57BL/6J** | |
|---|---|
| **Group A** | **Q1 vs. Q4<br>Adj. p-value** |
| WT/+ Chow | >0.9999 |
| WT/+ PLX5622 | <0.0001 |
| TG/+ Chow | >0.9999 |
| TG/+ PLX5622 | 0.0002 |
| One-way ANOVA<br>Quartile effect | F = 8.564<br>p-value = <0.0001 |

### S4K. PWK between Quartiles- ANOVA with Bonferroni post-hoc tests

| **PWK/PhJ** | |
|---|---|
| **Group A** | **Q1 vs. Q4<br>Adj. p-value** |
| WT/+ Chow | >0.9999 |
| WT/+ PLX5622 | >0.9999 |
| TG/+ Chow | >0.9999 |
| TG/+ PLX5622 | 0.6132 |
| One-way ANOVA<br>Quartile effect | F = 1.673<br>p-value = 0.1127 |

### Table S5- Associated Statistics for Tuft Dendrites & Spine Densities, related to Figure 3 and Figure S4

#### S5A. B6 Dendrites- ANOVA with Bonferroni post-hoc tests

| C57BL/6J | | Length | Branches |
|---|---|---|---|
| **Group A** | **Group B** | **Adj. p-value** | **Adj. p-value** |
| WT/+ Chow | WT/+ PLX5622 | 0.1012 | 0.1139 |
| WT/+ Chow | TG/+ Chow | >0.9999 | >0.9999 |
| WT/+ Chow | TG/+ PLX5622 | >0.9999 | >0.9999 |
| WT/+ PLX5622 | TG/+ Chow | 0.1962 | 0.1367 |
| WT/+ PLX5622 | TG/+ PLX5622 | 0.1949 | 0.5380 |
| TG/+ Chow | TG/+ PLX5622 | >0.9999 | >0.9999 |
| Two-way ANOVA (p-values): | | Treatment: 0.0578<br>Genotype: 0.0815<br>Interaction: 0.0784 | Treatment: 0.0326<br>Genotype: 0.1085<br>Interaction: 0.2338 |

#### S5B. PWK Dendrites- ANOVA with Bonferroni post-hoc tests

| PWK/PhJ | | Length | Branches |
|---|---|---|---|
| **Group A** | **Group B** | **Adj. p-value** | **Adj. p-value** |
| WT/+ Chow | WT/+ PLX5622 | >0.9999 | >0.9999 |
| WT/+ Chow | TG/+ Chow | >0.9999 | >0.9999 |
| WT/+ Chow | TG/+ PLX5622 | >0.9999 | >0.9999 |
| WT/+ PLX5622 | TG/+ Chow | >0.9999 | >0.9999 |
| WT/+ PLX5622 | TG/+ PLX5622 | >0.9999 | >0.9999 |
| TG/+ Chow | TG/+ PLX5622 | >0.9999 | >0.9999 |
| Two-way ANOVA (p-values): | | Treatment: 0.6461<br>Genotype: 0.9853<br>Interaction: 0.8992 | Treatment: 0.5874<br>Genotype: 0.6438<br>Interaction: 0.5468 |

#### S5C. Sholl Analysis- ANOVA

| | Dendrite length: Two-way ANOVA | | | | | |
|---|---|---|---|---|---|---|
| | C57BL/6J | | | PWK/PhJ | | |
| **Distance from bifurcation (um)** | **Treatment p-value** | **Genotype p-value** | **Interaction p-value** | **Treatment p-value** | **Genotype p-value** | **Interaction p-value** |
| 20 | 0.6848 | 0.3022 | 0.8257 | 0.8911 | 0.8219 | 0.9537 |
| 40 | 0.6356 | 0.6746 | 0.7734 | 0.7241 | 0.3215 | 0.8711 |
| 60 | 0.4821 | 0.8671 | 0.5285 | 0.8257 | 0.6272 | 0.7505 |
| 80 | 0.6596 | 0.6562 | 0.0786 | 0.3004 | 0.3683 | 0.1229 |
| 100 | 0.5167 | 0.5590 | 0.1081 | 0.3966 | 0.7228 | 0.1004 |
| 120 | 0.5682 | 0.9939 | 0.2387 | 0.8568 | 0.9452 | 0.3271 |
| ***140*** | 0.9322 | 0.6785 | 0.0057 | 0.7555 | 0.6157 | 0.7891 |
| ***160*** | 0.0864 | 0.8691 | 0.0181 | 0.2237 | 0.5677 | 0.9743 |
| 180 | 0.1091 | 0.3524 | 0.2096 | 0.5050 | 0.6169 | 0.6371 |
| 200 | 0.0511 | 0.3618 | 0.4097 | 0.6203 | 0.8432 | 0.8366 |
| ***220*** | 0.0317 | 0.8235 | 0.6480 | 0.1913 | 0.4503 | 0.5672 |
| 240 | 0.0724 | 0.4572 | 0.9554 | 0.5928 | 0.3514 | 0.9265 |
| 260 | 0.0643 | 0.1469 | 0.8299 | 0.5648 | 0.3673 | 0.6592 |
| ***280*** | 0.1217 | 0.0135 | 0.2804 | 0.3738 | 0.3319 | 0.1978 |
| ***300*** | 0.1442 | 0.0003 | 0.4607 | 0.6634 | 0.3322 | 0.4537 |
| ***320*** | 0.2804 | 0.0001 | 0.4122 | 0.8008 | 0.4458 | 0.7736 |
| ***340*** | 0.5166 | 0.0005 | 0.4344 | 0.8163 | 0.9289 | 0.8604 |
| ***360*** | 0.7212 | 0.0330 | 0.2931 | 0.7342 | 0.6903 | 0.9751 |
| ***380*** | 0.8943 | 0.0343 | 0.1795 | 0.6097 | 0.8302 | 0.3720 |
| 400 | 0.3782 | 0.1262 | 0.0899 | 0.4735 | 0.9156 | 0.2687 |
| 420 | 0.5219 | 0.1688 | 0.2269 | 0.7198 | 0.9214 | 0.2660 |
| 440 | 0.7926 | 0.7366 | 0.2028 | 0.8036 | 0.8761 | 0.2959 |
| 460 | 0.7043 | 0.7043 | 0.4073 | 0.5752 | 0.4044 | 0.2261 |

### S5D. B6 Sholl Analysis- Bonferroni post-hoc tests

| C57BL/6J | Bonferroni multiple comparison, adjusted p-value | | | | | |
|---|---|---|---|---|---|---|
| Distance from Bifurcation (um) | WT/+ Chow vs. WT/+ PLX5622 | WT/+ Chow vs. TG/+ Chow | WT/+ Chow vs. TG/+ PLX5622 | WT/+ PLX5622 vs. TG/+ Chow | WT/+ PLX5622 vs. TG/+ PLX5622 | TG/+ Chow vs. TG/+ PLX5622 |
| 140 | 0.0735 | 0.1907 | >0.9999 | >0.9999 | 0.1081 | 0.2421 |
| 160 | 0.0018 | 0.2535 | >0.9999 | 0.8152 | 0.3359 | >0.9999 |
| 220 | 0.0125 | >0.9999 | 0.0420 | 0.2424 | >0.9999 | 0.4826 |
| 280 | 0.2499 | 0.2424 | >0.9999 | 0.0019 | 0.0686 | >0.9999 |
| 300 | 0.2703 | 0.0261 | 0.2640 | 0.0001 | 0.0031 | >0.9999 |
| 320 | 0.9512 | 0.0846 | 0.2166 | 0.0042 | 0.0140 | >0.9999 |
| 340 | >0.9999 | 0.5135 | 0.5649 | 0.1472 | 0.1697 | >0.9999 |
| 360 | >0.9999 | >0.9999 | >0.9999 | 0.9265 | 0.5131 | >0.9999 |
| 380 | >0.9999 | >0.9999 | >0.9999 | >0.9999 | 0.8210 | >0.9999 |

### S5E. B6 Spine Density & Head Diameter- ANOVA with Bonferroni post-hoc tests

| C57BL/6J | | Density | Head Diameter |
|---|---|---|---|
| Group A | Group B | Tukey's post-hoc adj. p-value | Kolgorov-Smirnov p-value |
| WT/+ Chow | WT/+ PLX5622 | >0.9999 | <2.2e-16 |
| WT/+ Chow | TG/+ Chow | >0.9999 | 0.006598 |
| WT/+ Chow | TG/+ PLX5622 | >0.9999 | 0.149 |
| WT/+ PLX5622 | TG/+ Chow | 0.8848 | 2.243e-14 |
| WT/+ PLX5622 | TG/+ PLX5622 | >0.9999 | <2.2e-16 |
| TG/+ Chow | TG/+ PLX5622 | 0.3107 | 0.0004718 |
| Two-way ANOVA (p-values): | | Treatment: 0.0812<br>Genotype: 0.8348<br>Interaction: 0.2081 | |

### S5F. B6 Spine Density & Head Diameter- ANOVA with Bonferroni post-hoc tests

| PWK/PhJ | | Density | Head Diameter |
|---|---|---|---|
| Group A | Group B | Tukey's post-hoc adj. p-value | Kolgorov-Smirnov p-value |
| WT/+ Chow | WT/+ PLX5622 | >0.9999 | 0.08547 |
| WT/+ Chow | TG/+ Chow | 0.9246 | 0.000226 |
| WT/+ Chow | TG/+ PLX5622 | 0.0227 | 0.04984 |
| WT/+ PLX5622 | TG/+ Chow | 0.0354 | 0.01541 |
| WT/+ PLX5622 | TG/+ PLX5622 | 0.0002 | 0.06235 |
| TG/+ Chow | TG/+ PLX5622 | 0.7437 | 0.1749 |
| Two-way ANOVA (p-values): | | Treatment: 0.8784<br>Genotype: <0.0001<br>Interaction: 0.044 | |

### S5G. Comparisons to AT and ssEM data

| Heuer vs. Bloss Comparisons | | Density |
|---|---|---|
| Group A | Group B | Bonferroni post-hoc adj. p-value |
| Heuer B6 WT/+ Chow Confocal | Bloss AT | 0.9036 |
| Heuer B6 WT/+ Chow Confocal | Bloss EM | >0.9999 |
| Bloss AT | Bloss EM | >0.9999 |
| One-Way ANOVA: | | F = 0.7564<br>P = 0.4714 |

### S5H. B6 Spine head diameter within Quartile- ANOVA with Bonferroni post-hoc tests

| C57BL/6J | | Quartile (adj. p-value) | |
|---|---|---|---|
| **Group A** | **Group B** | **Q1** | **Q4** |
| WT/+ Chow | WT/+ PLX5622 | 0.0211 | 0.0980 |
| WT/+ Chow | TG/+ Chow | >0.9999 | >0.9999 |
| WT/+ Chow | TG/+ PLX5622 | >0.9999 | >0.9999 |
| WT/+ PLX5622 | TG/+ Chow | 0.0126 | >0.9999 |
| WT/+ PLX5622 | TG/+ PLX5622 | 0.2413 | 0.0122 |
| TG/+ Chow | TG/+ PLX5622 | >0.9999 | 0.7056 |
| Two-way ANOVA (p-value): | | Treatment: 0.0447<br>Genotype: 0.0652<br>Interaction: 0.1331 | Treatment: 0.8457<br>Genotype: 0.0062<br>Interaction: <0.0001 |

### S5I. B6 Spine head diameter within Quartile- ANOVA with Bonferroni post-hoc tests

| PWK/PhJ | | Quartile (adj. p-value) | |
|---|---|---|---|
| **Group A** | **Group B** | **Q1** | **Q4** |
| WT/+ Chow | WT/+ PLX5622 | >0.9999 | >0.9999 |
| WT/+ Chow | TG/+ Chow | >0.9999 | >0.9999 |
| WT/+ Chow | TG/+ PLX5622 | >0.9999 | 0.8056 |
| WT/+ PLX5622 | TG/+ Chow | >0.9999 | 0.1508 |
| WT/+ PLX5622 | TG/+ PLX5622 | >0.9999 | 0.0102 |
| TG/+ Chow | TG/+ PLX5622 | >0.9999 | >0.9999 |
| Two-way ANOVA (p-value): | | Treatment: 0.9781<br>Genotype: 0.5826<br>Interaction: 0.1207 | Treatment: 0.6366<br>Genotype: <0.0001<br>Interaction: 0.0511 |

### S5J. B6 Between Quartiles- ANOVA with Bonferroni post-hoc tests

| C57BL/6J | |
|---|---|
| **Group A** | **Q1 vs. Q4<br>Adj. p-value** |
| WT/+ Chow | >0.9999 |
| WT/+ PLX5622 | 0.0005 |
| TG/+ Chow | >0.9999 |
| TG/+ PLX5622 | 0.9558 |
| One-way ANOVA<br>Quartile effect | F = 4.419<br>p-value = <0.0001 |

### S5K. PWK Between Quartiles- ANOVA with Bonferroni post-hoc tests

| PWK/PhJ | |
|---|---|
| **Group A** | **Q1 vs. Q4<br>Adj. p-value** |
| WT/+ Chow | >0.9999 |
| WT/+ PLX5622 | >0.9999 |
| TG/+ Chow | >0.9999 |
| TG/+ PLX5622 | >0.9999 |
| One-way ANOVA<br>Quartile effect | F = 2.666<br>p-value = 0.0100 |

### Table S6- Associated Statistics for Oblique Dendrite Microglia Touch Analysis, related to Figure 4 and Figure S5

#### S6A. Touch vs. No Touch: Dendrite Count

| | C57BL/6J | | PWK/PhJ | |
|---|---|---|---|---|
| **Group** | **Microglia Touch (N)** | **Microglia NO Touch (N)** | **Microglia Touch (N)** | **Microglia NO Touch (N)** |
| WT/+ Chow | 42 | 25 | 26 | 29 |
| TG/+ Chow | 24 | 27 | 23 | 37 |
| WT/+ PLX5622 | 6 | 44 | 3 | 55 |
| TG/+ PLX5622 | 1 | 30 | 4 | 56 |

#### S6B. B6 Touch vs. No Touch: Spine Density & Head Diameter- non-parametric t-tests

| C57BL/6J | | Density | Head Diameter |
|---|---|---|---|
| **Group A** | **Group B** | **Two-tailed unpaired t-test p-value** | **Kolgorov-Smirnov p-value** |
| WT/+ Chow TOUCH | WT/+ Chow NO TOUCH | 0.0128 | 0.0003751 |
| TG/+ Chow TOUCH | TG/+ Chow NO TOUCH | 0.0144 | 0.0006093 |
| WT/+ Chow TOUCH | TG/+ CHOW TOUCH | NA | 0.09955 |
| WT/+ Chow NO TOUCH | TG/+ CHOW NO TOUCH | NA | 6.754e-9 |

#### S6C. PWK Touch vs. No Touch: Spine Density & Head Diameter- non-parametric t-tests

| PWK/PhJ | | Density | Head Diameter |
|---|---|---|---|
| **Group A** | **Group B** | **Two-tailed unpaired t-test p-value** | **Kolgorov-Smirnov p-value** |
| WT/+ Chow TOUCH | WT/+ Chow NO TOUCH | 0.1873 | 0.01979 |
| TG/+ Chow TOUCH | TG/+ Chow NO TOUCH | 0.1412 | 0.2076 |
| WT/+ Chow TOUCH | TG/+ CHOW TOUCH | NA | 0.1071 |
| WT/+ Chow NO TOUCH | TG/+ CHOW NO TOUCH | NA | 5.768e-7 |

#### S6D. B6 Proximal vs. Distal to Touch: Spine Density & Head Diameter

| C57BL/6J | | Density | Head Diameter |
|---|---|---|---|
| **Group A** | **Group B** | **Two-tailed Paired t-test p-value** | **Kolgorov-Smirnov p-value** |
| WT/+ Chow Proximal | WT/+ Chow Distal | 0.2911 | 0.6731 |
| TG/+ Chow Proximal | TG/+ Chow Distal | 0.3153 | 0.443 |

#### S6E. PWK Proximal vs. Distal to Touch: Spine Density & Head Diameter

| PWK/PhJ | | Density | Head Diameter |
|---|---|---|---|
| **Group A** | **Group B** | **Two-tailed Paired t-test p-value** | **Kolgorov-Smirnov p-value** |
| WT/+ Chow Proximal | WT/+ Chow Distal | 0.1056 | 0.07051 |
| TG/+ Chow Proximal | TG/+ Chow Distal | 0.5803 | 0.853 |

### Table S7- Associated Statistics for Tuft Dendrite Microglia Touch Analysis, related to Figure 4 and Figure S5

#### S7A. Touch vs. No Touch: Dendrite Count

| | C57BL/6J | | PWK/PhJ | |
|---|---|---|---|---|
| Group | Microglia Touch (N) | Microglia NO Touch (N) | Microglia Touch (N) | Microglia NO Touch (N) |
| WT/+ Chow | 46 | 17 | 17 | 37 |
| TG/+ Chow | 32 | 18 | 31 | 29 |
| WT/+ PLX5622 | 12 | 38 | 11 | 44 |
| TG/+ PLX5622 | 6 | 24 | 10 | 42 |

#### S7B. B6 Touch vs. No Touch: Spine Density & Head Diameter

| C57BL/6J | | Density | Head Diameter |
|---|---|---|---|
| Group A | Group B | Two-tailed Unpaired t-test p-value | Kolgorov-Smirnov p-value |
| WT/+ Chow TOUCH | WT/+ Chow NO TOUCH | 0.1061 | 0.0004951 |
| TG/+ Chow TOUCH | TG/+ Chow NO TOUCH | 0.0505 | 0.0002994 |
| WT/+ Chow TOUCH | TG/+ CHOW TOUCH | NA | 0.003892 |
| WT/+ Chow NO TOUCH | TG/+ CHOW NO TOUCH | NA | 3.043e-5 |

#### S7C. PWK Touch vs. No Touch: Spine Density & Head Diameter

| PWK/PhJ | | Density | Head Diameter |
|---|---|---|---|
| Group A | Group B | Two-tailed Unpaired t-test p-value | Kolgorov-Smirnov p-value |
| WT/+ Chow TOUCH | WT/+ Chow NO TOUCH | 0.5798 | 0.5203 |
| TG/+ Chow TOUCH | TG/+ Chow NO TOUCH | 0.2327 | 0.07733 |
| WT/+ Chow TOUCH | TG/+ CHOW TOUCH | NA | 0.4683 |
| WT/+ Chow NO TOUCH | TG/+ CHOW NO TOUCH | NA | 0.587 |

#### S7D. B6 Proximal vs. Distal to Touch: Spine Density & Head Diameter- nonparametric t-tests

| C57BL/6J | | Density | Head Diameter |
|---|---|---|---|
| Group A | Group B | Two-tailed Paired t-test p-value | Kolgorov-Smirnov p-value |
| WT/+ Chow Proximal | WT/+ Chow Distal | 0.0514 | 0.8362 |
| TG/+ Chow Proximal | TG/+ Chow Distal | 0.8609 | 0.06835 |

#### S7E. B6 Proximal vs. Distal to Touch: Spine Density & Head Diameter- nonparametric t-tests

| PWK/PhJ | | Density | Head Diameter |
|---|---|---|---|
| Group A | Group B | Two-tailed Paired t-test p-value | Kolgorov-Smirnov p-value |
| WT/+ Chow Proximal | WT/+ Chow Distal | 0.0395 | 0.2284 |
| TG/+ Chow Proximal | TG/+ Chow Distal | 0.4798 | 0.01178 |
